# Systematic characterization of genome editing in primary T cells reveals proximal genomic insertions and enables machine learning prediction of CRISPR-Cas9 DNA repair outcomes

**DOI:** 10.1101/404947

**Authors:** Ryan T. Leenay, Amirali Aghazadeh, Joseph Hiatt, David Tse, Judd F. Hultquist, Nevan Krogan, Zhenqin Wu, Alexander Marson, Andrew P. May, James Zou

## Abstract

The *Streptococcus pyogenes* Cas9 (SpCas9) nuclease has become a ubiquitous genome editing tool due to its ability to target almost any location in DNA and create a double-stranded break^1,2^. After DNA cleavage, the break is fixed with endogenous DNA repair machinery, either by non-templated mechanisms (e.g. non-homologous end joining (NHEJ) or microhomology-mediated end joining (MMEJ)), or homology directed repair (HDR) using a complementary template sequence^3,4^. Previous work has shown that the distribution of repair outcomes within a cell population is non-random and dependent on the targeted sequence, and only recent efforts have begun to investigate this further^5–11^. However, no systematic work to date has been validated in primary human cells^5,7^. Here, we report DNA repair outcomes from 1,521 unique genomic locations edited with SpCas9 ribonucleoprotein complexes (RNPs) in primary human CD4+ T cells isolated from multiple healthy blood donors. We used targeted deep sequencing to measure the frequency distribution of repair outcomes for each guide RNA and discovered distinct features that drive individual repair outcomes after SpCas9 cleavage. Predictive features were combined into a new machine learning model, CRI**SP**R **R**epair **OUT**come (SPROUT), that predicts the length and probability of nucleotide insertions and deletions with R^2^ greater than 0.5. Surprisingly, we also observed large insertions at more than 90% of targeted loci, albeit at a low frequency. The inserted sequences aligned to diverse regions in the genome, and are enriched for sequences that are physically proximal to the break site due to chromatin interactions. This suggests a new mechanism where sequences from three-dimensionally neighboring regions of the genome can be inserted during DNA repair after Cas9-induced DNA breaks. Together, these findings provide powerful new predictive tools for Cas9-dependent genome editing and reveal new outcomes that can result from genome editing in primary T cells.

## Main

A wide range of therapeutic applications based on genome engineering of human hematopoietic cells are under active development. Primary T cells present a notably promising cell type for therapeutic genome editing, as they can be engineered efficiently *ex vivo* and transferred adoptively to patients^12^. Despite this, we still lack detailed information about the genomic outcomes of Cas9-dependent editing in primary human cells. Here, we use targeted sequencing to measure DNA repair outcomes at 1,656 unique genomic locations targeting 559 genes (three guides per gene, with positive and negative/non-targeting controls) in primary CD4+ T cells isolated from 18 human donors. Primary CD4+ T cells were isolated from human blood donors and expanded as described previously^13^. Guide RNAs were combined with SpCas9 to assemble RNPs and electroporated into T cells^14,15^. After 6 days of recovery and expansion, DNA was isolated from cells electroporated with each RNP, and a 180-260 base pair (bp) region around each site was PCR amplified and sequenced (Fig. 1A).

**Figure 1.**
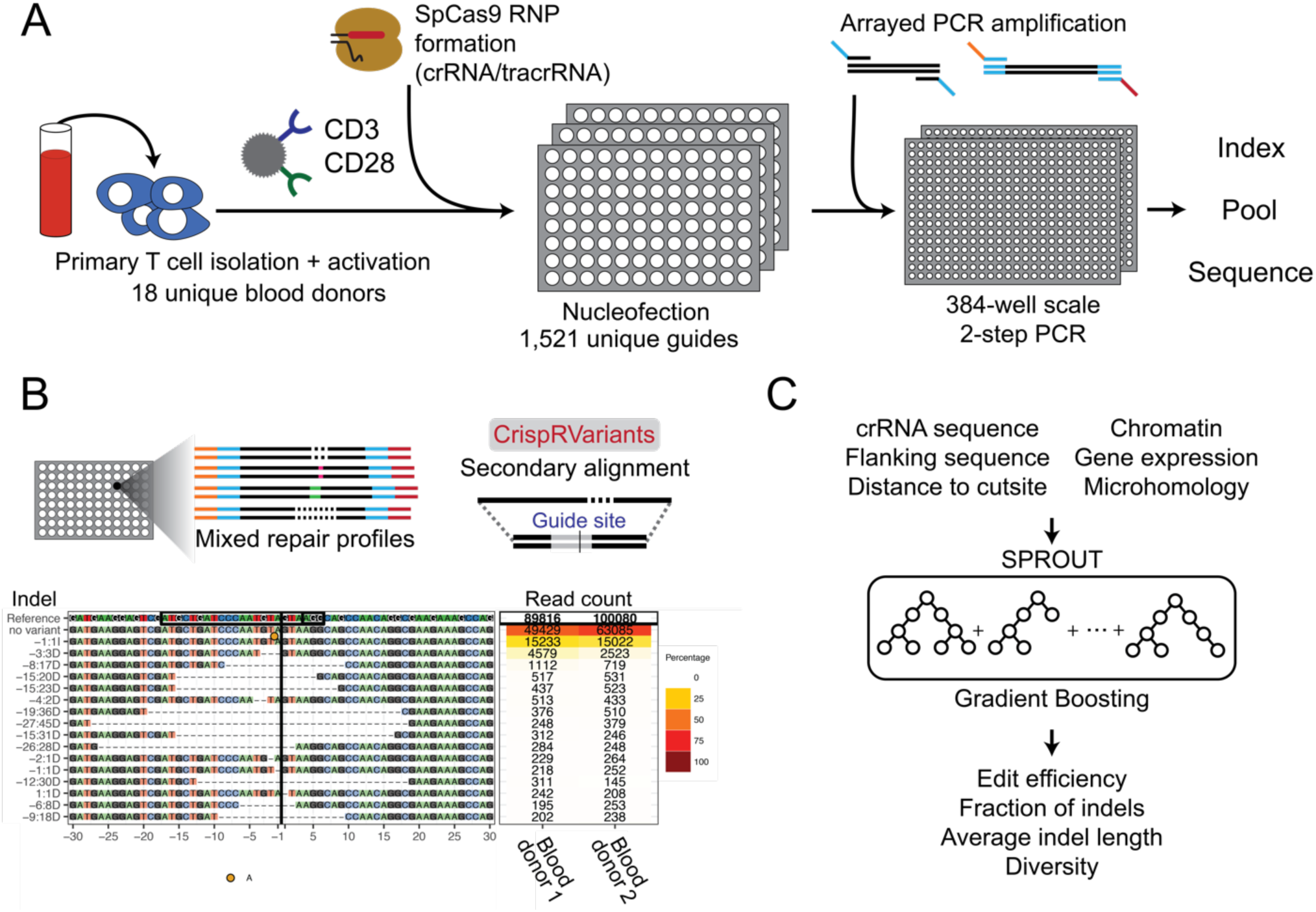
Overview of the method. **(A)** Primary T cells were isolated, activated, and electroporated with Cas9/crRNA/tracrRNA RNPs in 96-well plates. After 6 days of expansion, genomic DNA was isolated from each well, amplified and sequenced. **(B)** The CrispRVariants R package^16^ was used to quantify each SpCas9 RNP knockout. An example alignment is plotted here, with quantification shown for two blood donors. Each site has this same unique plot, all of which can be found on figshare. **(C)** A gradient boosting machine learning algorithm was trained to predict multiple DNA repair outcomes given the guide RNA sequence, chromatin factors, and gene expression levels.

We quantified the distribution of repair outcomes at each target site from the generated amplicon library using CrispRVariants^16^ (Fig. 1B). Target sites with fewer than 1,000 insertion or deletion (indel)-containing reads were removed from further analysis to ensure repair outcomes were measured accurately. There were 1,521 unique target sites from 549 genes that passed this filtering, with an average read depth of 57,555; 1,361 of these sites were replicated in two or more donors (Supplementary Fig. 1). We focused our analysis on these 1,521 high quality target sites.

In total, 31% of reads contained deletions centered around the cut site with an average deletion length of 13 bps. We also found that 20% of the reads had insertions at the cut site, and 95% of these insertions were of exactly one nucleotide. Only 0.008% of reads contained both an insertion and deletion. Moreover, fewer than 1% of the reads contained single nucleotide variation (SNV) but no indel, some of which may be attributable to sequencing error, and we chose to focus our analysis on reads containing at least one insertion or deletion.

DNA repair outcomes were distinct at each site, similar to previous observations from immortalized cell lines^5^. There was an average of 98 discrete repair outcomes per site that were observed at a frequency greater than 1 in 1000 reads, and different target sites were highly variable in the proportion and length distribution of insertions and deletions. The standard deviation of the ratio of insertions to deletions was 0.43 across all cut sites (Supplementary Fig. 2-4) with 25% and 75% quantiles equal to 0.08 and 0.41, respectively. Repair outcomes from sites were compared systematically by converting the top 20 indels for each site into binary bit strings and calculating Jaccard similarity coefficients (Fig. 2A, 2B). Three control sites, targeting CXCR4, LEDGF and CDK9, were tested in cells derived from each of the donors. The repair outcomes from each target site were very similar between donors, but very different between different target sites (Fig. 2A). Comparisons of repair outcomes between all sites showed that replicate editing experiments from individual target sites were significantly more similar to each other than to outcomes from different sites (Fig. 2B, Supplementary Fig. 5). Overall, these data demonstrate that DNA repair outcomes in primary T cells are highly variable but non-random and largely consistent between cells from different donors.

**Figure 2.**
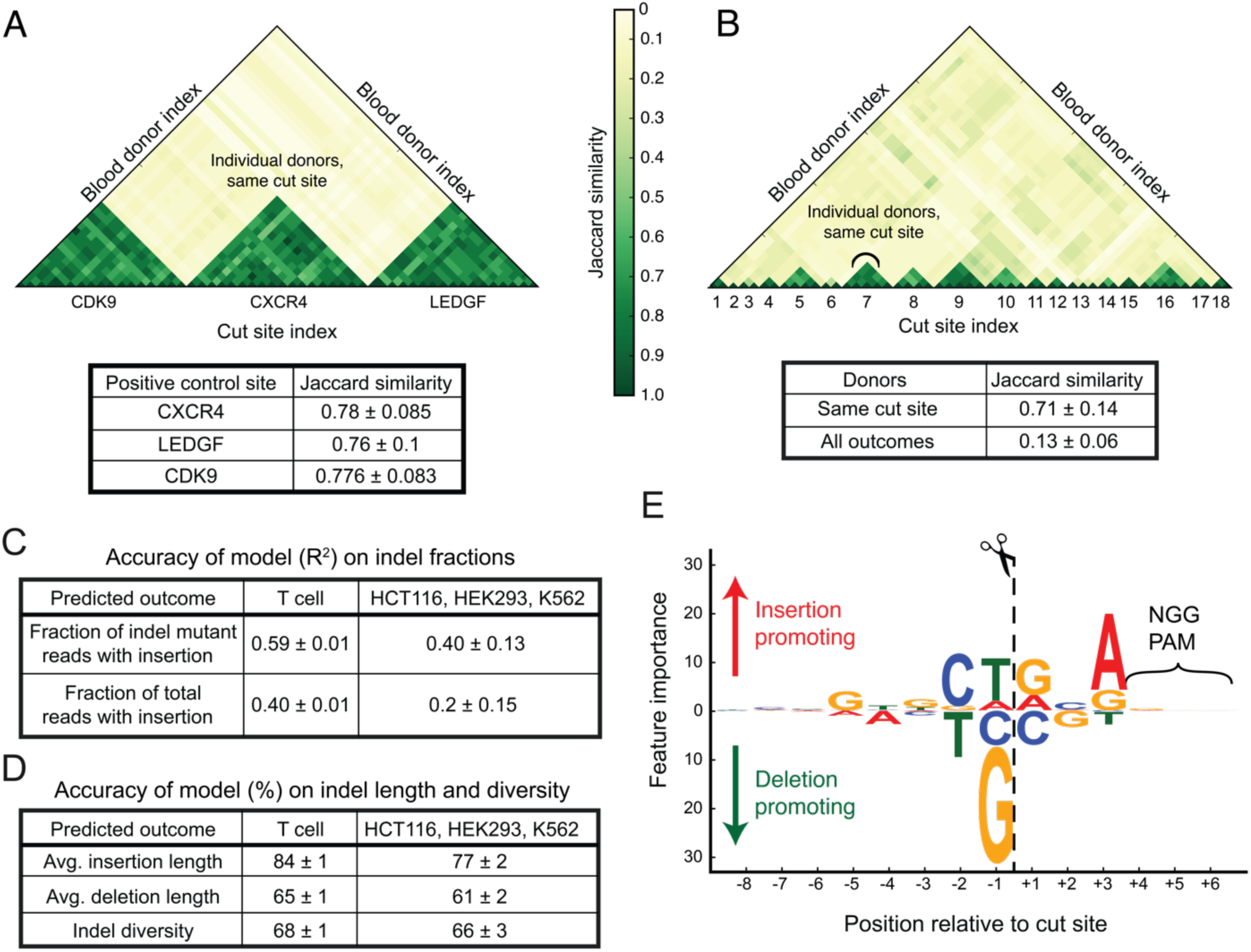
SPROUT predicts DNA repair outcomes. **(A)** The DNA repair outcomes resulting from RNP activity in T cells derived from different blood donors were compared for CDK9, CXCR4, and LEDGF control guides analyzing the top 20 indels at each site. These guides were used in every blood donor. Jaccard similarity is calculated for each guide site across donors. **(B)** Jaccard similarity of DNA repair outcomes for 18 randomly chosen guides, again using the top 20 indels. Jaccard coefficients are plotted comparing outcomes from different guide RNAs and between blood donors. **(C)** The trained model was used to predict DNA repair indel fractions in a hold-out (un-seen) portion of the T cell dataset. The model was also evaluated on previously published data^5^ obtained from immortalized cell lines to test generalization performance for other cell types and experimental conditions. **(D)** Accuracy of the trained model in predicting the average insertion and deletion length and indel diversity on both T cells and previously published data^5^. **(E)** The importance that SPROUT assigns to nucleotides at each position relative to the cut site. Larger text indicates that the presence of a particular nucleotide at a position has greater importance in determining the likelihood of insertion versus deletion.

We hypothesized that the variation in repair outcomes across cut sites was largely due to sequence variation near the cut site^5–7^ and would enable prediction based on sequence. To test this, we developed a machine learning model, SPROUT, to predict SpCas9 repair outcomes (Fig. 1C). The model takes as input the 20 nucleotides of the spacer sequence plus the PAM. At each target site, the model predicted the fraction of indel mutant reads with an insertion (Fig. 2C) and deletion (1 - fraction of insertions) and the average length of insertions and deletions (Fig. 2D). We trained and evaluated SPROUT using five-fold cross validation. On an independent set of 304 target sites in primary T cells, SPROUT was able to accurately predict the fraction of indel mutant reads with an insertion (R^2^ = 0.59) and the fraction of total reads with an insertion (R^2^ = 0.40, Fig. 2C).

The prediction signal was primarily localized in the three nucleotides immediately to the left and right of the cut site. When SPROUT was trained using just these six nucleotides, it predicted the fraction of indel-containing reads and total reads with insertions with an R^2^ of 0.56 and 0.38, respectively. Incorporating additional features such as local chromatin status and gene expression did not improve the predictive power for SpCas9 repair outcomes. SPROUT also predicted the edit efficiency with an accuracy of R^2^ = 0.23, where about 52% of the prediction power was due to the guide sequence alone (with R^2^ = 0.12) and the remaining power was derived from chromatin features and gene expression (with R^2^ = 0.11). These results were compared to other similar algorithms, showing improved performance in all cell types and especially in T cells^17,18,19,20^ (Supplementary Fig. 6).

We then investigated whether SPROUT could correctly select which SpCas9 target site in a gene was the most likely to have an enrichment of insertions in order to create a tool for *in silico* guide design. For each of the 532 genes with multiple guides, we used SPROUT’s prediction to rank the target sites by their predicted fraction of indel mutant reads with an insertion. For 73% of the genes, SPROUT correctly chose the top sgRNA, and for 60% of genes it correctly predicted the complete ranking of all the candidate guides by their insertion proportion, significantly above random guessing (Supplementary Fig. 7, p < 10^−154^).

We also evaluated which DNA sequence features affected DNA repair outcomes^5,7^, and we discovered that the −1 position (immediately to the 5’ end of the cleavage site) was the most influential (Fig. 2E and Supplementary Fig. 8). This is consistent with previous observations where this nucleotide is duplicated at many cut sites, which has been suggested to be the result of repair of single-base overhangs generated by Cas9^6^. The presence of a G or C nucleotide at this position decreased the insertion proportion: 7% and 10% of indel mutant reads were insertions, respectively. Comparatively, the presence of A or T nucleotide at this position increased this proportion to 23% and 26%, respectively. The +3 position is also important in determining the proportion of outcomes as insertions or deletions (Supplementary Fig. 8B). A or G nucleotides at this position increase the insertion proportion to 25% and 23% respectively, compared with 16% and 15% for C and T. The presence of homopolymers (a run of two or more identical nucleotides) adjacent to the cut site increased the proportion of deletions (p < 0.02). For example, targets with G homopolymers abutting the cut site have deletions in 92% of the indel mutant reads, compared to 77% deletions when there is no homopolymer at the cut site (Supplementary Fig. 9), which could be a reflection of microhomology mediated end joining^4^.

Next, we assessed the robustness of the algorithm to sequence- and cell-specific features by using the SPROUT model trained on the T cell data to predict SpCas9 repair outcomes in other human cell types. We re-analyzed published targeted sequencing data from 96 unique target sites tested in HEK293, K562, and HCT116 cells^5^ (Supplementary Fig. 10). These 96 targets were distinct from the 1,521 sites that were used to train SPROUT, and hence constitute new test data. SPROUT achieved an accuracy of R^2^ = 0.40 in predicting the fraction of indel mutant reads with an insertion and an R^2^ = 0.23 in predicting the fraction of total reads with an insertion. The relatively high cross-cell-type performance of SPROUT further suggests that the primary factor influencing the repair outcomes after SpCas9 cleavage within dividing cells is the nucleotide sequence context near the cut site. However, the lower accuracy compared with the T cell data strongly suggest that cell type and experimental conditions may have additional influence on the distribution of repair outcomes, and that cell-type-specific models may be required for accurate prediction.

In the process of developing the SPROUT model to predict predominant insertions (Supplementary Fig. 11-12), we determined that the majority of insertions were less than 3 nucleotides in length (Supplementary Fig. 4). However, for 99% of the target sites, we also observed insertions that were at least three nucleotides in length, corresponding to 1.7% of all insertion-containing reads. Within this subset there were a prevalent class of insertions, over 25 nucleotides in length, present at low frequency per target site (Fig. 3A and Supplementary Fig. 13). These long insertions were found within 1,406 unique sites, corresponding to 92% of assessed cut sites (Fig. 3B). 40% of these long insertions aligned to the human genome (Fig. 3B); the remaining long insertions did not align with the default settings of BLAST. Averaged across sites, 0.07% of indel-containing reads (46 reads per site) were aligned long insertions. These aligned long insertions were not enriched for repetitive sequences nor for SINE motifs (Supplementary Materials). Only 36% aligned to regions within 1kb of the cut site (Supplementary Fig. 14) while the remaining 64% of long inserted sequences aligned to a different chromosome than the target site. The origins of the long insertions, i.e. the donor sites, were enriched for enhancers and transcriptionally active genomic regions (Supplementary Fig. 15). T cells from different individuals edited at the same cut site were significantly more likely to share the same long insertions compared to random pairs of cut sites (p < 10^−39^). The same cut sites from different individuals were also more likely to have large insertions that come from similar regions in the genome compared to different cut sites from the same individual (Fig. 3C, p < 10^−101^). Moreover, different cut sites within the same gene were more likely to share closer long insertions compared to cut sites from different genes (p < 10^−95^).

**Figure 3.**
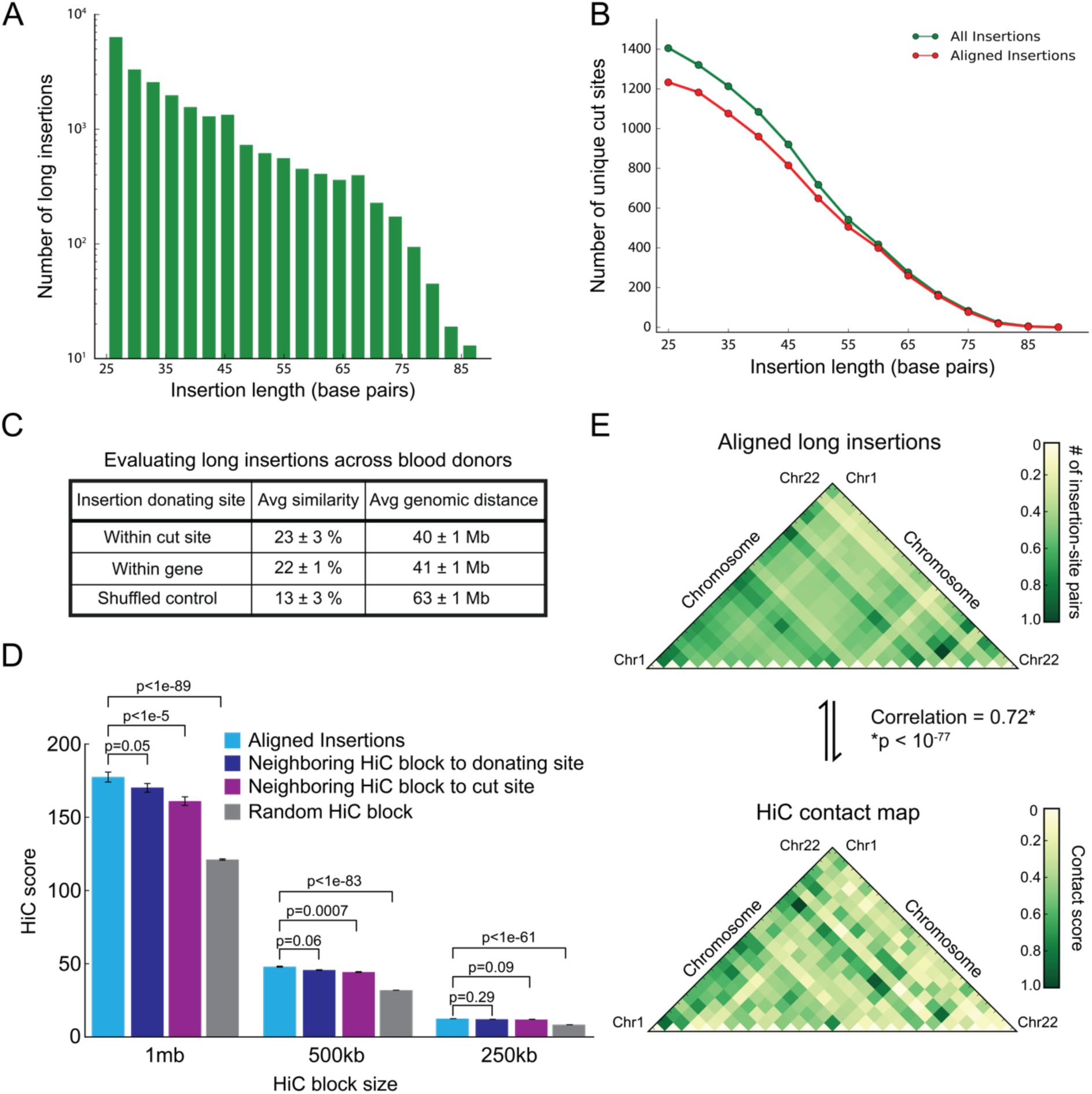
Genomic DNA sequences from sites in physical proximity can be inserted at SpCas9 cut sites. **(A)** All insertions longer than or equal to 25 bases were identified and plotted. **(B)** Number of cut sites with at least one (aligned) long insertion was plotted against the insertion length. **(C)** Average similarity of the insertion location within the same cut site and gene was compared across donors. This was also performed on a shuffled set of insertions as a control. Average genomic distance quantifies the distance between the donor sites of the long insertions that originate from the same chromosome. **(D)** HiC chromosome contact maps were directly compared to the aligned long insertions. **(E)** Quantification of the HiC contact data to the long insertions. Neighboring blocks as well as a randomly selected block were used as controls. The HiC block size was also varied.

Together, these results suggested that the physical location of the cut site affects the type of insertions that occur during DNA repair (Fig. 3C). To test this hypothesis, we used HiC data^21^ to quantify the physical proximity between the cut site and the location of origin of the inserted sequence, which we denote as the donor site. Cut sites had more HiC contacts with donor sites than they did with regions that are adjacent to the donor (p = 0.05) or with random regions (Fig. 3D, p < 10^−89^). Conversely, donor sites were also more likely to have HiC contacts with the cut site than with regions adjacent to the cut site (p < 10^−5^). Due to the resolution limit of the HiC data, we found that the signal was more strongly enriched when we considered regions of sizes 500 kb and 1 mb (Fig. 3D). At the inter-chromosomal level, the likelihood of having long insertion exchanges between two chromosomes, normalized for chromosome length, was highly correlated with the proximity between the chromosomes, as measured via HiC (Fig. 3E, p < 10^−77^). Furthermore, we verified that insertions enriched for proximal genomic sequences were also found after editing in HEK293 and K562 cells (Supplementary Fig. 16,17). The enrichment for HiC contacts remained significant, although the smaller number of target sites in those data reduced the strength of the signal. Together, this analysis suggests that insertion of proximal genomic sequences occurs at low frequency in multiple cell types, and could be a general consequence of SpCas9-induced double-stranded DNA breaks.

## Discussion

Here, we publish the largest analysis to date of DNA repair outcomes resulting from SpCas9 editing in primary human CD4+ T cells. We used this dataset to develop SPROUT, a prediction algorithm to determine repair outcomes after a SpCas9 double-stranded DNA break. SPROUT has significantly enhanced prediction capacity compared to previous models in primary human T cells and, to a lesser extent, in transformed cell lines. It is able to rank guide RNA sequences by proportion of insertions and deletions and has revealed key elements of the nucleotide sequence that drive repair outcomes. Consistent with previous results^5–7^, we find that the nucleotides adjacent to the cut site are the major driver of NHEJ repair outcomes and that the presence of homopolymers increase the fraction of deletions. We also reveal an unanticipated role for the PAM-adjacent nucleotide in promoting insertions as a repair outcome. This nucleotide on the non-template strand packs against the end of the PAM duplex at the point the two strands diverge^22^. It is tempting to speculate that interaction of purine nucleobases with the PAM duplex will result in a different conformation of the non-template strand to pyrimidine nucleobases, resulting in a shift in likelihood towards generating single-base staggered cuts through the RuvC domain. Similarly, the −1 nucleotide is critical for positioning the non-template strand for cleavage by the RuvC domain and is involved in extensive interactions with both Cas9 protein and other nucleotides from the strand. Future structural and biochemical studies with different guide and target sequences will be required to illuminate this in detail. It important to note that while many of the features of repair outcomes were shared between primary T cells and transformed cell lines, SPROUT was more accurate at predicting repair outcomes for primary T cells. This suggests that there may be cell-type specific effects, and that any models targeted toward prediction of outcomes will require training on the specific cell-type of interest to achieve high accuracy. This may be of particular importance in post-mitotic cells, where influences of the cell cycle on DNA-repair will be more pronounced than in rapidly dividing populations.

These experiments also reveal that the repair outcomes from the majority of target sites contain low frequency long DNA insertions that originate from other parts of the genome. A substantial fraction of these insertions align to regions of the genome that are physically proximal to the SpCas9 target site. Our analyses show that the long insertions preferentially originate from enhancers and transcriptionally active regions of the genome, and that they are not repetitive elements or retrotransposons. Recent reports have indicated that cells may undergo significant genomic rearrangements in response to SpCas9 cleavage^8,23^, although these should be interpreted cautiously given the cell types used. Short-range targeted PCR based sequencing is unable to detect many of these reported larger rearrangements, and these would have been missed in our analysis. However, despite this lack of sensitivity, the prevalence of the proximal genomic insertions in our data set suggests that some fraction of cells may use accessible regions of the genome to repair DNA using non-canonical mechanisms. Long-range PCR followed by long read sequencing or high coverage genome sequencing could provide additional data on complex repair outcomes, and perhaps reveal additional DNA sources^8,24^. However, long-range PCR and long read sequencing are inherently biased towards shorter fragments, and improved methods will be required for accurate quantification of the true occurrence rate of these insertion events. Given the therapeutic applications of T cells and other primary cells, it would be prudent to investigate further into the mechanisms and prevalence of these, and other larger rearrangements, during genome editing.

## Data availability

The SPROUT software is publicly available at https://github.com/amirmohan/SPROUT.git. All raw data and analyses are openly available through SRA (BioProject: PRJNA486372) and figshare, respectively.

## Methods

### T cell Editing

Lyophilized crRNA and tracrRNA (Dharmacon) was resuspended at a concentration of 160 µM in 10 mM Tris-HCL (7.4 pH) with 150 mM KCl. Cas9 ribonucleoproteins (RNPs) were made as previously described by combining 5µL of 160µM crRNA with 5µL of 160µM tracrRNA for 30 min at 37°C, followed by incubation of this 80µM gRNA product with 10µL of 40µM Cas9 (UC Macrolab) to form RNPs at 20µM^25^. Five 3.5µL aliquots were frozen in lo-bind 96-well V-bottom plates (E&K Scientific) at −80° C until used. All crRNA guide sequences were designed by Dharmacon for gene knockout (Hiatt *et al*, in review).

T cell editing was conducted according to published protocols^14^. Briefly, peripheral blood mononuclear cells (PBMC) were isolated from whole blood (numeric donors, under a protocol approved by UCSF Committee on Human Research, CHR #13-11950) or de-identified residuals from leukoreduction chambers after Trima Apheresis (alphabetic donors, from Blood Centers of the Pacific) from healthy human donors by Ficoll centrifugation with SepMate tubes (STEMCELL, per manufacturer’s instructions). CD4+ T cells were then isolated from PBMCs with magnetic negative selection (STEMCELL), cultured at 1 million cells/mL in complete RPMI (RPMI-1640 with 20 IU/mL IL2, 10% FBS, 50 µg/mL Pen/Strep and 5mM HEPES) and activated with plate-bound anti-CD3 (OKT3) and anti-CD28 (CD28.2) antibodies.

After three days of culture on stimulating antibodies at 37°C / 5% CO_2_, cells were resuspended and counted before editing. Approximately 3.5 x 10^5^ cells were edited per blood donor per guide. Immediately before electroporation, cells were centrifuged at 400xg for 5 minutes, supernatant was aspirated, and the pellet resuspended in 20 µL of room-temperature Lonza electroporation buffer P3 (Lonza). The cell suspension was then gently mixed with thawed RNP and carefully aliquoted into 96-well electroporation cuvette for nucleofection with the 4D 96 well shuttle unit (Lonza) using code EH-115. Immediately after electroporation, 80 µL of pre-warmed media without IL2 were added to each well and cells were allowed to rest for at least one hour in a 37°C cell culture incubator. Subsequently cells were moved to 96-well flat-bottomed culture plates pre-filled with 100 µL warm complete media with IL2 at 40 IU/mL (for a final concentration of 20 IU/mL) and anti-CD3/anti-CD2/anti-CD28 beads (T cell Activation and Stimulation Kit, Miltenyi Biotec) or anti-CD3/anti-CD28 dynabeads (ThermoFisher) at 1:1 bead:to:cell ratio.

Cells were then cultured at 37°C / 5% CO_2_ in a dark cell culture incubator for a further 6 days, and were supplemented with IL2-containing complete media on days 3 and 5 of culture. On day 6 of culture, one eighth of each culture, approximately 35 µL, was reserved for genomic DNA analysis by 1:1 mixing with QuickExtract buffer (EpiCentre) in a 96-well plate, sealing carefully with foil and heating to 65°C for 20 min followed by heating to 98°C for 5 minutes on a thermocycler. Genomic DNA extracts were stored at −20°C until use.

### PCR amplification of cut sites

PCR primers were designed using an in-house Python wrapper around Primer3 (github.com/czbiohub/Primer3Wrapper)^26^. Primers were designed to amplify a 180 to 260 nucleotide region, ensuring that the cut site was at least 50 nucleotides from the end of each primer, as well as 15 nucleotides from the center of the read to ensure there was enough sequence to accurately quantify larger indels. Sequencing adapters (Forward: 5’-CTCTTTCCCTACACGACGCTCTTCCGATCT-3’ and Reverse 5’-CTGGAGTTCAGACGTGTGCTCTTCCGATCT-3’) were appended to the designed primers, and a homodimer and heterodimer filter was applied to ensure no secondary structure existed between primers. Sites were amplified using between 4,000 and 10,000 genomic copies, 0.5 µM of each primer, and Q5 hot start high-fidelity 2x master mix (NEB). PCR was performed using the standard protocol: 98°C for 30 seconds; then 35 cycles of 98°C for 10 seconds, 60°C for 30 seconds, and 72°C for 30 seconds; followed by a final extension at 72°C for 2 minutes (NEB). Samples were diluted 1:100 and individually indexed in a second, 12-cycle PCR using index primers containing Illumina sequencing adapters and 8 base barcodes, under the same conditions as the first PCR. After the second PCR, indexed samples were pooled and purified using a 0.7x SPRIselect purification and sequenced on an Illumina NextSeq 500.

### Repair outcome pre-processing pipeline

Fastq sequencing files were first merged using FLASH^27^, then subjected to adapter and quality trimming with trimmomatic^28^. These merged reads were then initially aligned to the hg38 genomic contig using bwa mem^29^, creating individual .bam files. Each sample was individually analyzed using the CrispRVariants bioconductor package in R^16^, which performs a secondary alignment and quantifies each unique insertion and deletion per sequencing read. Repair outcomes were then further parsed using embedded CrispRVariants packages to quantify individual DNA repair outcomes, the insertion sequences, mutation efficiencies, and SNVs. Sites where the total number of reads was less than 1,000 were considered dropouts and filtered from all analysis. The average number of reads per site, after filtering, is approximately 59,000.

### T cell data summary

This study involved 3,989 DNA repair profiles from T cells isolated from 18 patients. These outcomes targeted 1,521 unique sites within 549 genes in the human genome. The RNP knockouts were repeated on average 2 times, each across unique primary T cells from different blood donors (Supplementary Fig. 1). The repair outcomes were averaged over the repeats across the patients, and DNA repair outcome data from each target site has been deposited on figshare.

### HCT116, HEK293, and K562 data summary

Published sequencing data^7^ from three other cell types (HEK293, K562, and HCT116; BioProject PRJNA326019) were analyzed according to the same procedure as the T cell data and used for validation of the machine learning model. The dataset we used from the manuscript comprised the RNP knockouts, after 48 hours, from 96 unique cut sites on the human genome. Each knockout was biologically repeated three times (Supplementary Fig. 10), and DNA repair outcome data from each target site has been deposited on figshare.

### SNVs within the dataset

Single nucleotide variations (SNVs) are rarely observed in SpCas9-based repair outcomes. On average, only 1.04% of our reads at a given cut site show one or more SNVs, with the 25% to 75% quantiles equal to 0.45% and 0.75%, respectively.

### Jaccard similarity

We computed the Jaccard similarity by analyzing the top 20 DNA repair outcomes. This analysis measures the ratio of the number of shared repair outcomes between the two sets over the total number of repairs in both sets together.

### Training the machine learning model

We used five-fold cross-validation to train SPROUT. We randomly split the unique cut sites in T cells (a total of 1,521) into 5 folds and trained SPROUT on four of the five folds. We then tested the performance of SPROUT on the remaining unseen fifth fold (304 cut sites). We repeated the random data split procedure 10 times and report the average and standard deviation of the prediction performance over the 10 random repeats. We evaluated the prediction performance of regression tasks, i.e., predicting the fraction of total or indel mutant reads with insertion or deletion and the edit efficiency, using the coefficient of determination (R^2^). We also evaluated the prediction performance of classification tasks, i.e., predicting if the average insertion or deletion length or the diversity is larger/smaller than the median of the distribution, using the accuracy of the classifier. A naive (or random) guess would be 50% accurate in predicting the correct output labels.

For the models evaluated on three other cell types (HCT116, HEK293, and K562), we trained SPROUT on the full T cell data (1,521 cut sites) and tested the performance of the model on the other cell types. For the classification tasks, we used the median of the cell type distributions to set the threshold.

### Fraction of insertions and average insertion length

We trained SPROUT to predict the fraction of reads with an insertion using both the number of reads with an insertion divided by the number of reads with insertion or deletion (fraction of indel mutant reads), and the number of reads with an insertion divided by the total number of reads, (fraction of total reads). We also trained SPROUT to predict if the expected insertion length is more or less than a threshold of the average insertion (approximately a single base). To determine SPROUT’s accuracy, a classifier was used based on the median size of the insertion (1.11 bp for T cells, 1.14 bp for HCT116/HEK293/K562s). The accuracy of this classifier was 84% and 77% for the two classes of cells tested, respectively. Note that random naive guess achieves 50% accuracy on this task (34% and 27% accuracy gain).

### Fraction of deletions and average deletion length

We replicated the method for insertions to predict deletion outcome. Due to the increased variance in deletion length compared to insertion length (standard deviation of 5.0 bp compared to 0.4 bp), the prediction was more difficult (Supplementary Fig. 4). SPROUT predicted the fraction of deletion mutant reads with an R^2^ = 0.59. Note that since the fraction of indel mutant reads with an insertion and deletion adds to one, the prediction performance of these two outcomes are similar. Our model predicted the fraction of total reads with a deletion with R^2^ = 0.10, significantly lower than the insertion prediction (R^2^ = 0.40, Fig. 2). The model did not retain accuracy on the immortalized human cell lines (average R^2^ = 0.05).

On T cells SPROUT predicted (using the previously described classification method) if the average deletion length is less or more than the median of 12.19 bps with an accuracy of 65% (15% accuracy gain over a random guess). We validated the model trained on T cells on other cell types, i.e., HCT116, HEK293, and K562; the median of the expected insertion length on other cell types was 11.76 bps. The model trained on T cells predicted if the average deletion length is less or more than the median with an accuracy of 61% (11% accuracy gain) in these three cell lines.

### Homopolymer analysis

To analyze the effect of homopolymers on indel formation, we fit a linear regression model based on spacer (+PAM) input sequence, whose output predicted the fraction of mutant reads containing a deletion. Four binary indicators measured the presence (or absence) of homopolymers. To obtain the binary indicator variables, we assess if the nucleotides adjacent or crossing the cut site repeat for more or equal to two times. Each of the four indicators proved to be significant in predicting deletions (p < 0.02). The length of homopolymers is also positively correlated with increased deletion proportion (p < 0.02).

### Predicting the diversity of outcomes

We use the entropy of the repair outcome distribution as a metric to quantify the site-specific repair diversity. This entropy measured the distribution of the repair outcomes – whether they were more diverse and widespread, or singular and enriched – again based on the input guide and PAM sequence. The model classified outcomes into two classes of high and low diversity, which correspond to having an entropy of less than or larger than the median (3.66) of entropies across repair outcomes. This classifier had an accuracy of 68% on T cells. This generalized to the immortalized lines with 70% accuracy (median of 3.05).

### Nucleotide features extracted from the machine learning model

To measure the importance of individual features in the gradient boosting model, the information gain concept was used. The information gain associated to a feature measures the decrease in entropy after a dataset is split based on that particular feature. A higher information gain corresponds to a more predictive feature. We also determined the influence of each feature (enrichment or depletion) from the sign of the coefficients of a linear regression model trained on the data. Note that the algorithm was completely blind to the actual location of the cut site. Additionally, the feature importance for nucleotides (e.g., ‘G’) showed an alternating pattern. We speculate that one reason for the enrichment of alternating pattern for an insertion outcome and thus depletion for a deletion outcome is the homopolymer effect. It has been observed that homopolymers – the repetition of one base creating long runs of the same nucleotide – favor a deletion outcome (Supplementary Fig. 9)^4,5^.

### Ranking guides based on a desired repair outcome

We evaluated SPROUT in ranking the guides based on fitness to produce a desired repair outcome. Two outputs were used to train the regression: the fractions of indel reads, and the fractions of total reads. After training on 400 genes, the model was used to predict the fraction of insertions and deletion of a hold-out set of guides targeting 149 different genes. We assessed the ranking performance of the guides on only the genes that have more than one guide in our datasets (142 test genes out of 149 genes total). The guides were then ranked within each gene based on the insertion and deletion fractions, and the rank correlation between the observed result and predicted ranking was evaluated.

The performance was measured using Kendall’s tau ranking coefficient and the percentage of completely correct predictions. Kendall’s tau ranking coefficient measures the difference between the observed result and the predicted rankings. The Kendall’s tau coefficient is a ranking measure between −1 and 1, where 1 indicates that rankings match exactly, 0 means that there is no ranking correlation, and −1 means that there is complete reverse ranking correlation. We tabulated the results in Supplementary Fig. 7 both in ranking the guides in hold-out genes from T cells and guides from the three other validation cell types (HCT116, HEK293, and K562).

### Edit efficiency prediction

We compared the performance of previously trained models for edit efficiency^17,18,30^ on the edit efficiency from the T cells repair data (Supplementary Fig. 6). These models showed no power (R^2^ < 0.01) in predicting the edit efficiency in T cells, HCT116, HEK293, or K562 (R^2^ < 0.01). This lack of power suggests that the published predictors are specific to the particular delivery methods and readouts used in these papers. SPROUT, in contrast, predicted the edit efficiency of T cells with R^2^ = 0.23. However, this did not carry over to other cell types.

### Predicting short inserted sequences

We trained SPROUT to predict the type of the nucleotide that is inserted during an insertion event of length one. We first attempted to predict if the inserted sequences are more likely to be A/T or C/G in a single-nucleotide insertion event (Supplementary Fig. 12). Of the total dataset, 1245 of the guides were more likely to have A/T insertion and the remaining 267 sites were more likely to have C/G insertion. Since this was an unbalanced dataset, we used the more proper F1 measure, which computes the harmonic mean of precision (TP / (TP + FP)) and recall (TP / (TP + FN)), where TP is the true positive, FN is the false negative, and FP is the false positive. Our model predicted the “dual” insertion type (A/T or G/C) with 96% accuracy (F1 = 0.87).

SPROUT was also trained to predict the insertion type among the four distinct classes A, C, G, and T, again given a single nucleotide insertion event. SPROUT predicted the inserted nucleotide with a 49% accuracy. Naive guessing strategy of the most frequent class achieved an accuracy of 43% (6% accuracy gain). Validation on the other cell types showed a prediction accuracy of 77% while the naive guessing strategy was also 77% accurate (no accuracy gain).

### Extracting and aligning long insertion data from the repair outcomes

To obtain the insertion data, the repair outcomes of all 1,521 cut sites were parsed and reads with an inserted sequence of length at least 25 bp were selected, totaling 22,495 unique insertions which centered on the cleavage site (Supplementary Fig. 11). All insertions were aligned to the human genome with the BLAST algorithm (blastn command, https://blast.ncbi.nlm.nih.gov/Blast.cgi) under default conditions and input parameters. For the cases with more than one alignment, the site with the highest alignment score was selected. A total of 8,946 unique insertions aligned to the human genome (Supplementary Fig. 13). 36% of the long insertions aligned to the same chromosome as that of the cleavage site (intra-chromosomal map) and the remaining insertions (64%) aligned to other chromosomes (inter-chromosomal map).

### HiC contact maps database

The HiC contact map of GM12878 primary cell lines (the most T cell-like line with data in public inter-chromosomal HiC databases) was used to evaluate the HiC contact scores in T cells. The contact maps are available at https://www.ncbi.nlm.nih.gov/geo/query/acc.cgi?acc=GSE63525^21^. HiC contact maps from various resolutions (250 kb, 500 kb, 1 mb) were used. The raw observed data (chrXX_YY_ZZb.RAWobserved) with the MAPQGE30 format was used. For the immortalized K562 line, the same GM12878 contact map was used. The HEK293 immortalized line was analyzed using the NHEK contact map [GEO accession number: GSE63525]. The chromosomal level contact maps were obtained by adding the pairwise scores between the blocks with chromosomes.

### HiC contact maps analysis

We employed a statistical analysis test to test the correlation between the HiC maps and the aligned insertions using different resolutions: both chromosomal level and genomic block level. For the chromosomal level, the average HiC map between chromosome pairs^21^ was compared to the mapped insertions (Fig. 3E), skipping the sex chromosomes because sex was masked in our T cell donors. This interaction was normalized by the length of the chromosomes, and the resulting correlation (Pearson correlation of 0.72, p < 10^−77^) suggested raw chromosome to chromosome contact correlated with the aligned insertions. To analyze genomic blocks, we collected and averaged the values of HiC scores and compared the values to three controls. The first control averaged the HiC contact scores on the neighboring HiC block to the selected blocks from the insertion donating site side (see Fig. 3A for visualization). The second control averaged the HiC scores on the neighboring HiC block to the selected blocks from the cut sit side. The third control averaged the HiC scores on random blocks within the same chromosome pairs that the cut happened and insertion aligned to. The results spanned different block sizes ranging from 250kb to 1mb.

### Repetition of long insertions across donors in the same cut sites and genes

We also quantified the similarity of the inter-chromosomal insertions (larger than 25bp) across the same cut sites (or genes). We analyzed the similarity between the aligned chromosomes, given a common cleavage site, and compared them with random control. To measure this, we defined S^ij^ = {c^ij^_1_,c^ij^_2_,c^ij^_3_,…,c^ij^_Nij_}, where the chromosome set is of cleavage site i, donor was j, and N_ij_ was defined as the unique long insertions. Thus, c^ij^_k_ denoted the chromosome index of the k^th^ insertion donating site, corresponding to donor j at cut site i. We then quantified the similarity of long insertion outcomes of donors j = {1,2,3,…,M^i^} at cut site i by finding the fraction of shared elements between sets S^ij^ and S^ij’^ for every pair of donors (j,j’). This was compared to a random control comprised of two random cut sites or genes from shuffled, distinct donors. The analysis shows that the long insertion outcomes which are observed on the same cut site are significantly (p < 10^−39^) closer to each other (23% similarity) compared to the random control case (13% similarity). The same observation was true for the long insertion outcomes observed on the same gene (22% similarity and p < 10^−17^). The same method was used to quantify across the biological replicates from published immortalized cell line data^5^.

We also quantified the chromosomal similarity of long insertions across donors, measured by the location of the aligned insertion. The average distance between inserted sequences was calculated and compared to a random control baseline (40 Mb vs 63 Mb average distance, respectively). Similar results were obtained when evaluating aligned long insertions obtained from cleavage sites within the same gene. This same method again was used to evaluate long insertions from previously published sequencing data^5^ (Supplementary Fig. 16).

### Insertion repeat element analysis

To evaluate if the aligned long insertions were enriched for repetitive sequences and for retrotransposons, we performed a computational search for common features. We calculated the frequency of k-mers, and the entropy of these k-mer frequencies. As an example, SINE elements which often have runs of (CA)_n_ and (CT)_n_^31^, corresponding to a low entropy. The average entropy of k-mers was calculated for each aligned long insertion and compared to a random shuffling of the nucleotides within an insertion. The average entropy of aligned insertions (3.18 ± 0.25) did not show a significant difference from random shuffled sequences (3.24 ± 0.24) when we analyzed the 3mer frequency. We also searched for consensus motifs of SINE (TGGCNNAGTGGN and GGTTCGANNCC) and only found the motifs in 4 aligned long insertions.

## Acknowledgments

This work was supported by the Chan Zuckerberg Biohub. J.Z. was supported by a Chan-Zuckerberg Investigator grant and by National Science Foundation grant CRII 1657155. A.M. was supported by NIH/NIDA Avenir New Innovator Award (DP2DA042423, A.M.), NIH/NIGMS funding for the HIV Accessory & Regulatory Complexes (HARC) Center (P50 GM082250, A.M. and N.J.K.), and a gift from Jake Aronov. A.M. holds a Career Award for Medical Scientists from the Burroughs Wellcome Fund and is an investigator at the Chan Zuckerberg BioHub. A. A. was supported by National Institute of Health grant 7R01HG008164-04 and the Stanford data science initiative. We would like to thank Norma Neff and Rene Sit for assistance collecting sequence data and Anna Sellas for lab support. This work builds on the work of the entire CRISPR and genome editing communities and we thank our friends and colleagues there for robust discussions along the path to this manuscript.

## Conflict of interests

A.M. is a co-founder of Spotlight Therapeutics. A.M. has served as an advisor to Juno Therapeutics and is a member of the scientific advisory board at PACT Pharma. The Marson laboratory has received sponsored research support from Juno Therapeutics, Epinomics, Sanofi and a gift from Gilead.

---

**Supplementary Figure 1.**
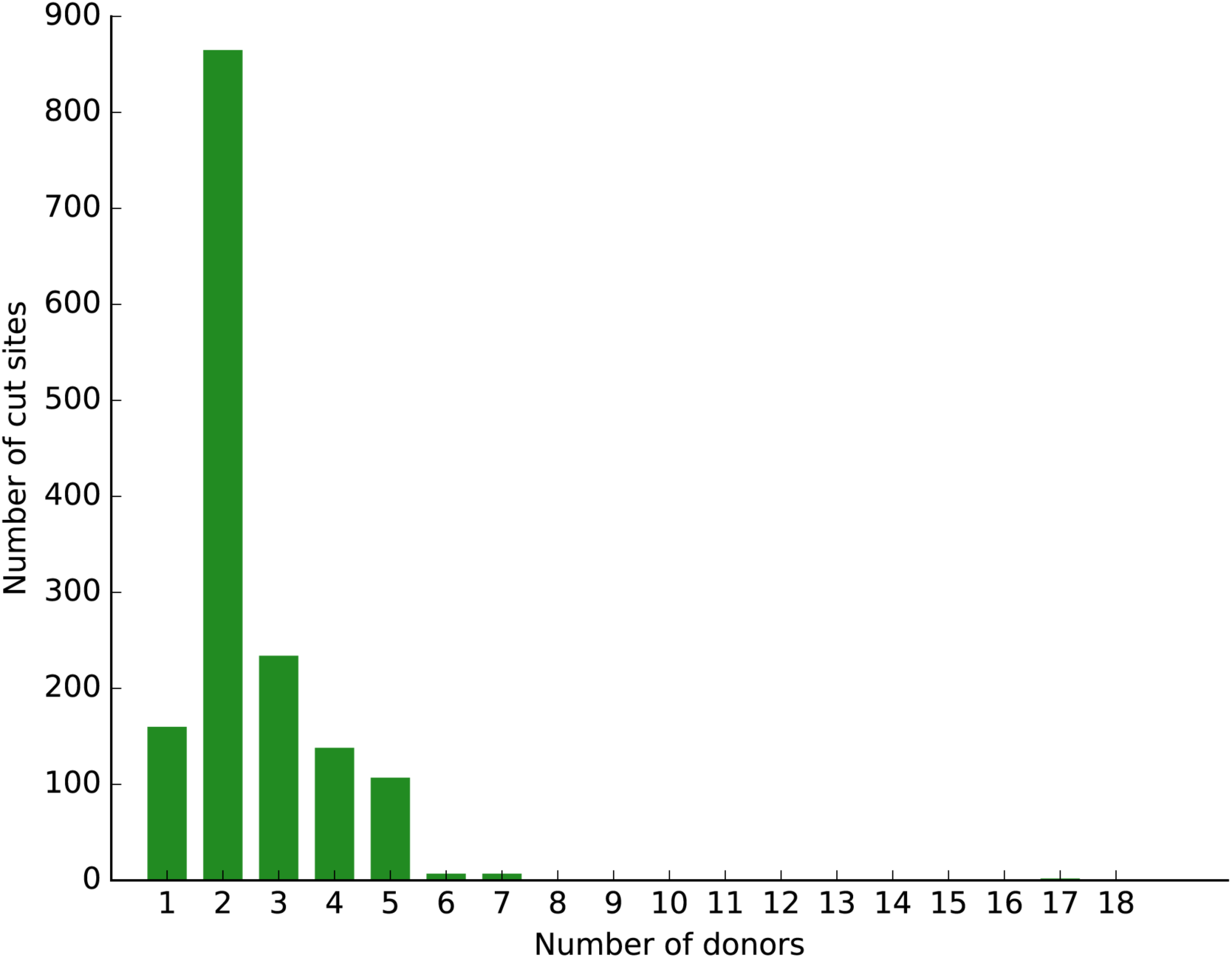
Histogram showing the distribution of unique blood donors per SpCas9 cut site in the T cell data.

**Supplementary Figure 2.**
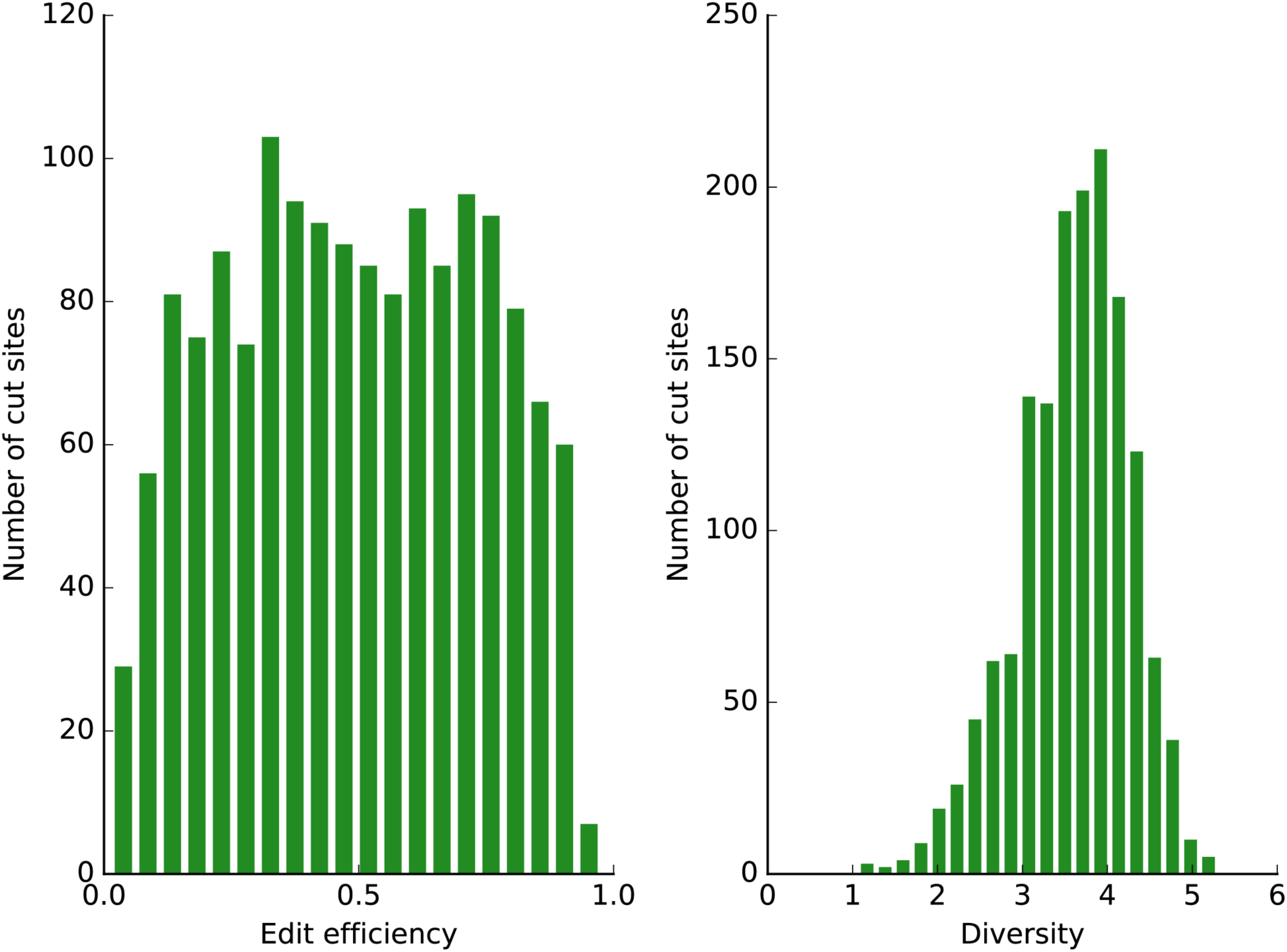
Distribution of the edit efficiency (left) and indel diversity (right) of the repair outcomes in T cells. We use the entropy of the distribution of the reads over the indel types as a metric to quantify the diversity of the repair outcomes. If there is exactly one repair outcome in all of the reads, then the entropy is 0. Higher entropy means that the repair outcomes are more diverse.

**Supplementary Figure 3.**
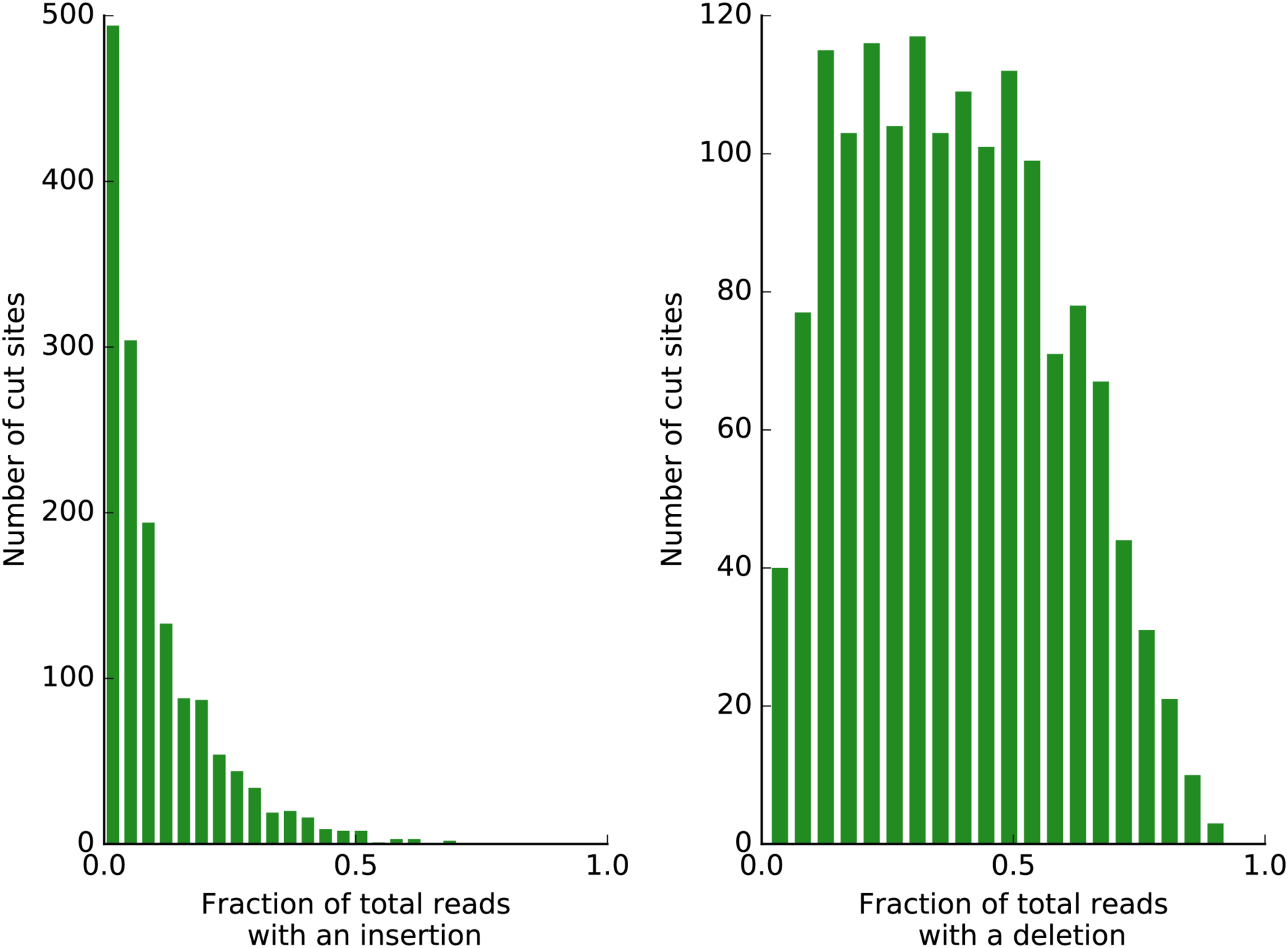
Distribution of the fraction of total reads with an insertion (left) and deletion (right) in T cells.

**Supplementary Figure 4.**
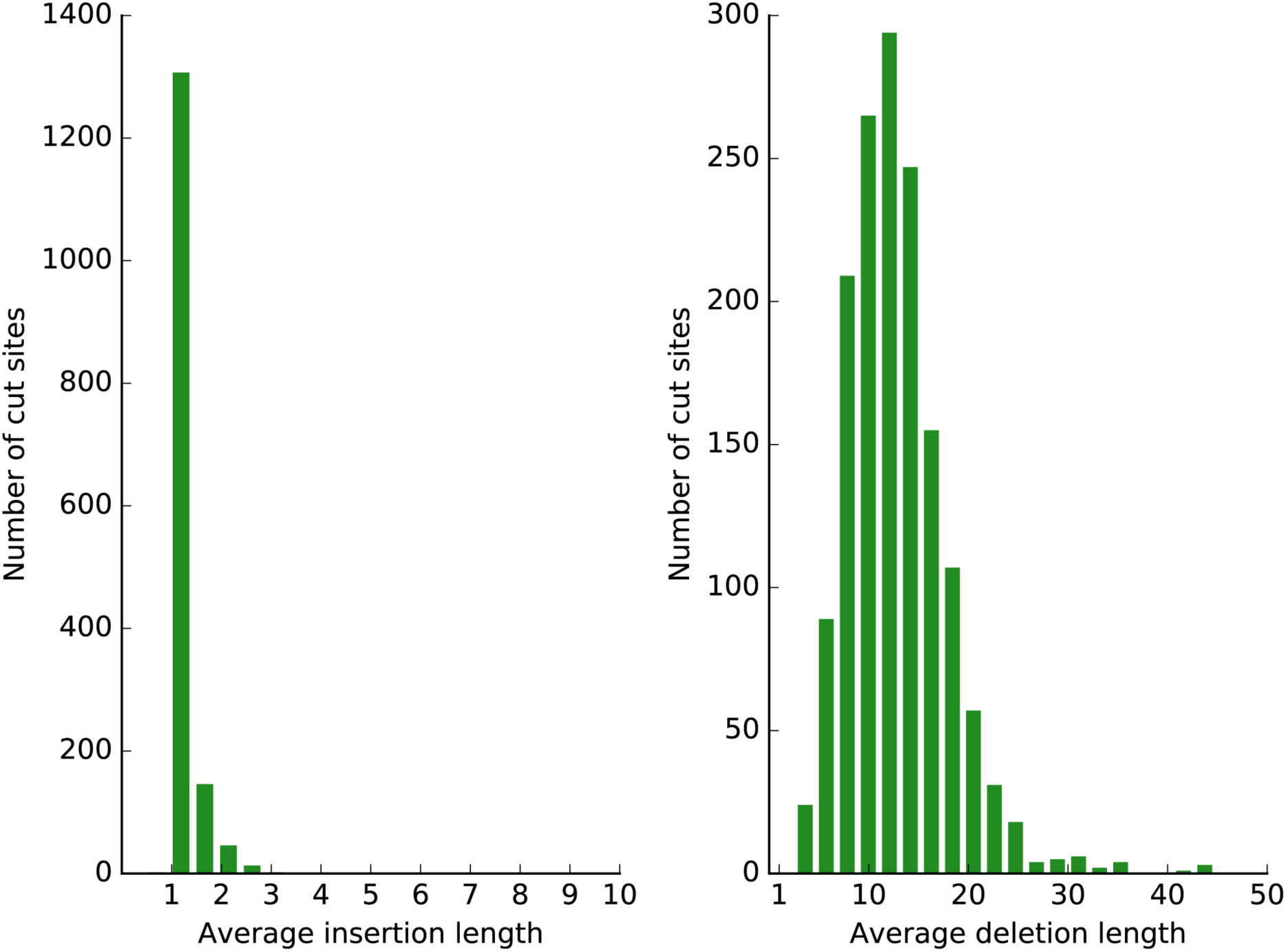
Distribution of the average insertion length given an insertion (left) and average deletion length given a deletion (right) in the repair outcomes of T cells.

**Supplementary Figure 5.**
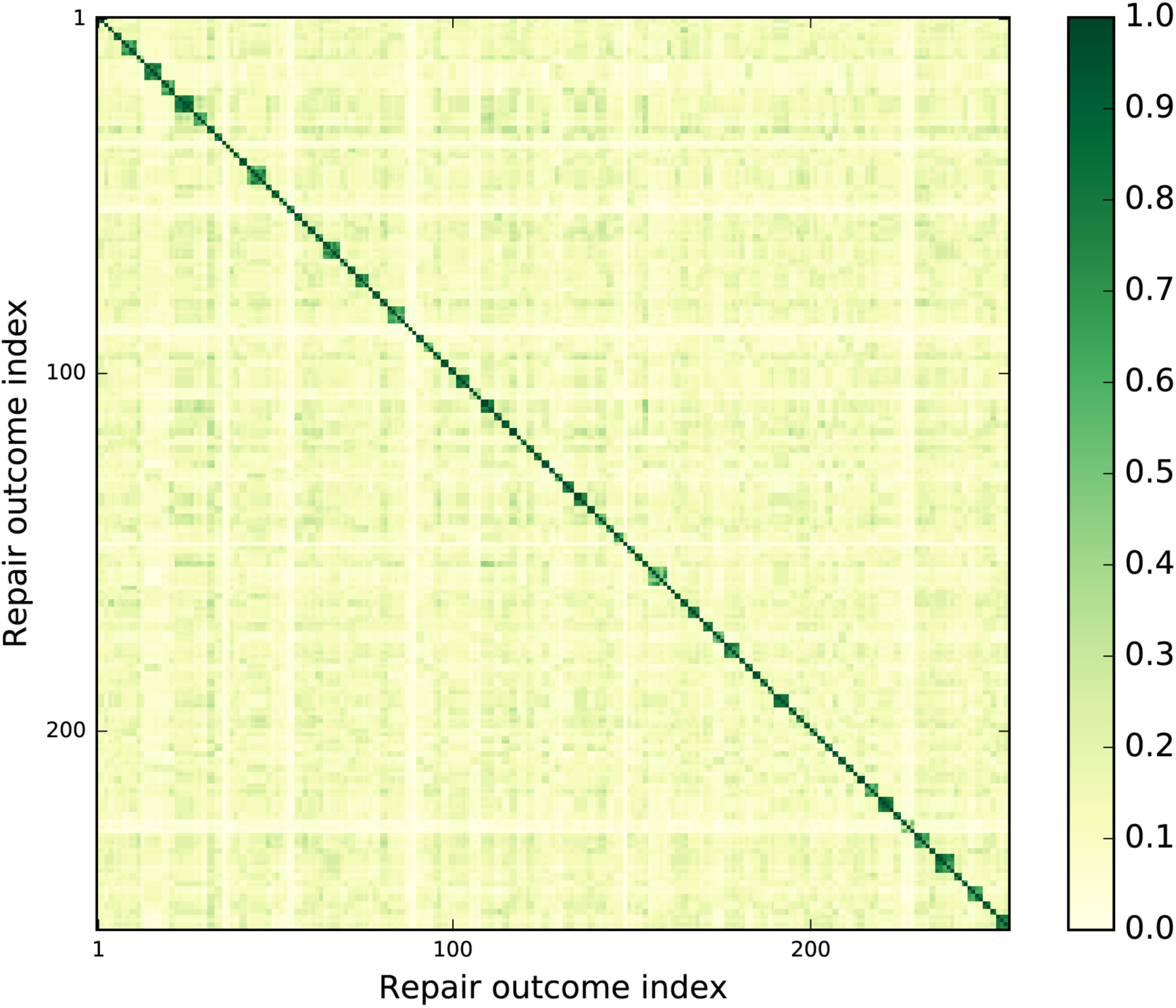
The Jaccard similarity between the top 20 indels of 250 (of a total of 3,989) randomly selected SpCas9 targeted sequencing experiments in T cells. Experiments performed on cells from different individuals at the same cut site are placed next to each other in the heatmap. These biological replicates show greater Jaccard similarity in repair outcomes compared to outcomes at distinct cut sites, as can be seen in the blocks along the diagonal.

**Supplementary Figure 6.**
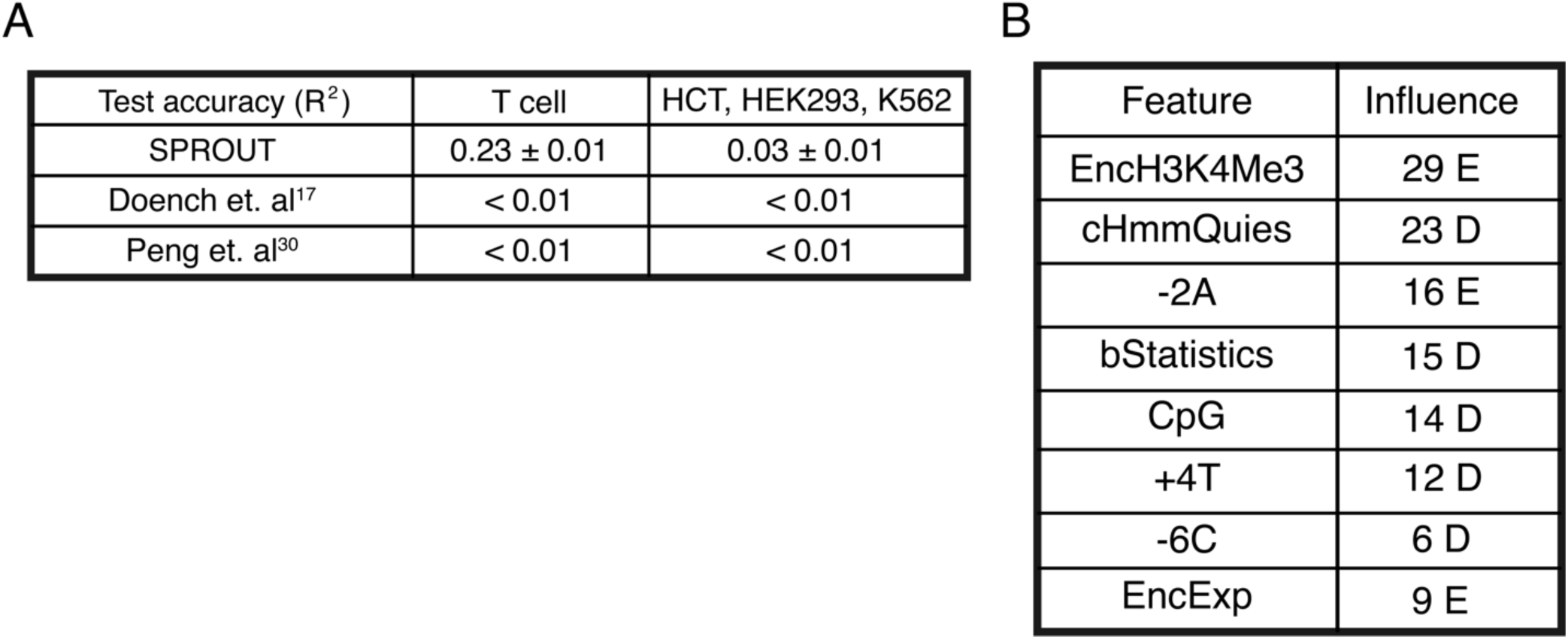
Performance of SPROUT in predicting editing efficiency. **(A)** Comparison of SPROUT machine learning model to two other published models^18,30^. SPROUT was trained on a random subset of T cell sites and the table indicates its accuracy on the hold-out T cell targets as well as on three other cell types on which it was not trained. **(B)** The eight most important features (ranked by information gain) of SPROUT for predicting edit efficiency. E indicates that presence of the feature corresponds to increasing the number of insertions relative to deletions, and D indicates the features corresponds to decreasing the number of insertions relative to deletions. See http://cadd.gs.washington.edu/static/ReleaseNotes_CADD_v1.2.pdf for more details on the extracted features from the ENCODE project.

**Supplementary Figure 7.**
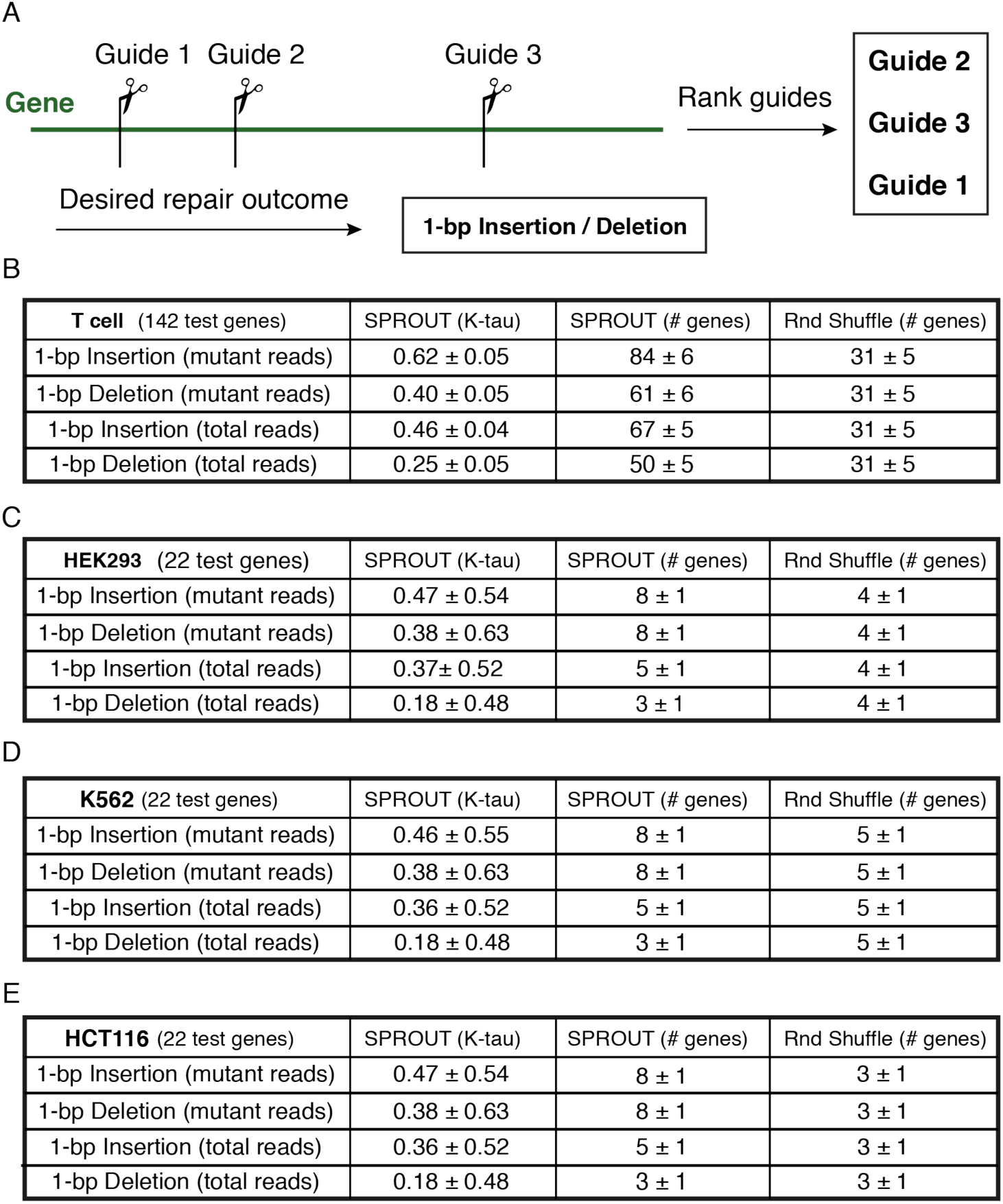
SPROUT’s performance in ranking guides within a gene based predicted repair outcome. **(A)** Schematic of the guide ranking experiment. Assuming a gene with three potential guides (Guide 1, Guide 2, Guide 3), SPROUT ranks the guides based on likelihood to produce a single nucleotide insertion (or deletion). In this illustration the algorithm predicts that Guide 2 produces the most number of reads with 1-bp insertion/deletion. **(B)** Guide ranking performance on T cells. The algorithm was trained on 435 genes and tested on the remaining 108 genes. Kendal tau (between [-1,1]) measures the rank correlation (higher is better and zero indicates no correlation) and “# genes” indicates the number of genes for which SPROUT predicted exactly the correct ranking across all the guides. **(C)** Guide ranking performance on HEK293. **(D)** Guide ranking performance on K562. **(E)** Guide ranking performance on HCT116. For parts (C,D,E) the model was trained on all T cell genes and tested on 28 genes from these other cell types.

**Supplementary Figure 8.**
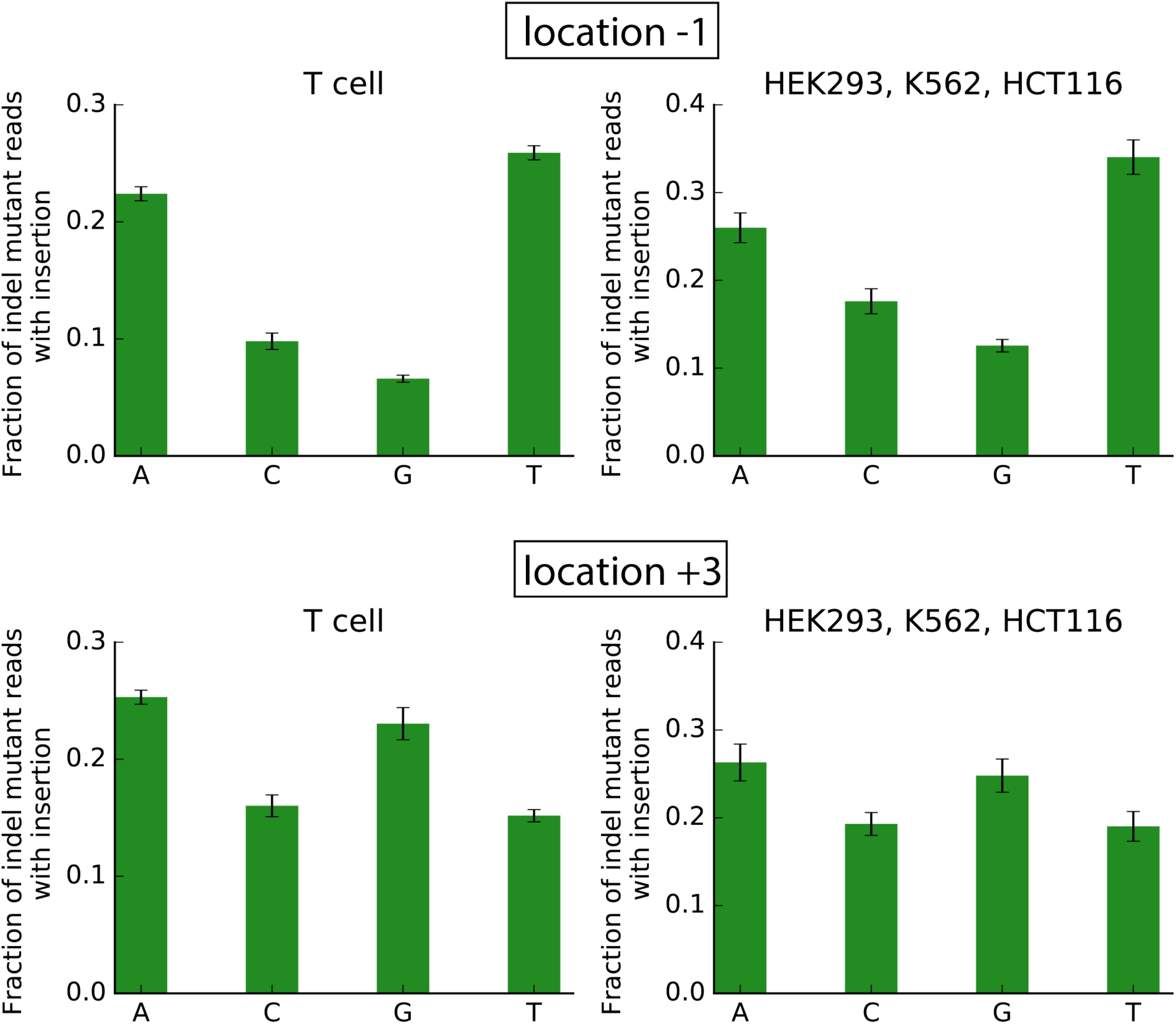
(A) Average fraction of indel mutant reads with insertion in target sites grouped by their nucleotide type at location −1 (adjacent to the cut site from the 5’ side). Presence of C or G at location −1 is significantly correlated with higher deletion proportion (p < 0.004) and presence of A or T is significantly correlated with higher insertion proportion (p < 10^−6^) consistently across all cell types. We show the results for T cells (left) and the aggregate results for HEK293, K562 and HCT116 (right). **(B)** Average fraction of indel mutant reads with insertion conditioned on the nucleotide at position +3 (the last nucleotide before e.g. 5’ of the PAM sequence). The presence of A at +3 is correlated with higher fraction of insertions. Error bars represent the standard error of the mean (SEM). The analyses here differ from and complement Fig 2E. The SPROUT importance scores of 2E captures the nonlinear model’s overall prediction as to the impact of each nucleotide and position. The figures here ignore the effects of other positions and plots the conditional insertion fractions. Even though the methods are different, both the feature importance scores and the conditional fractions give consistent biological findings.

**Supplementary Figure 9.**
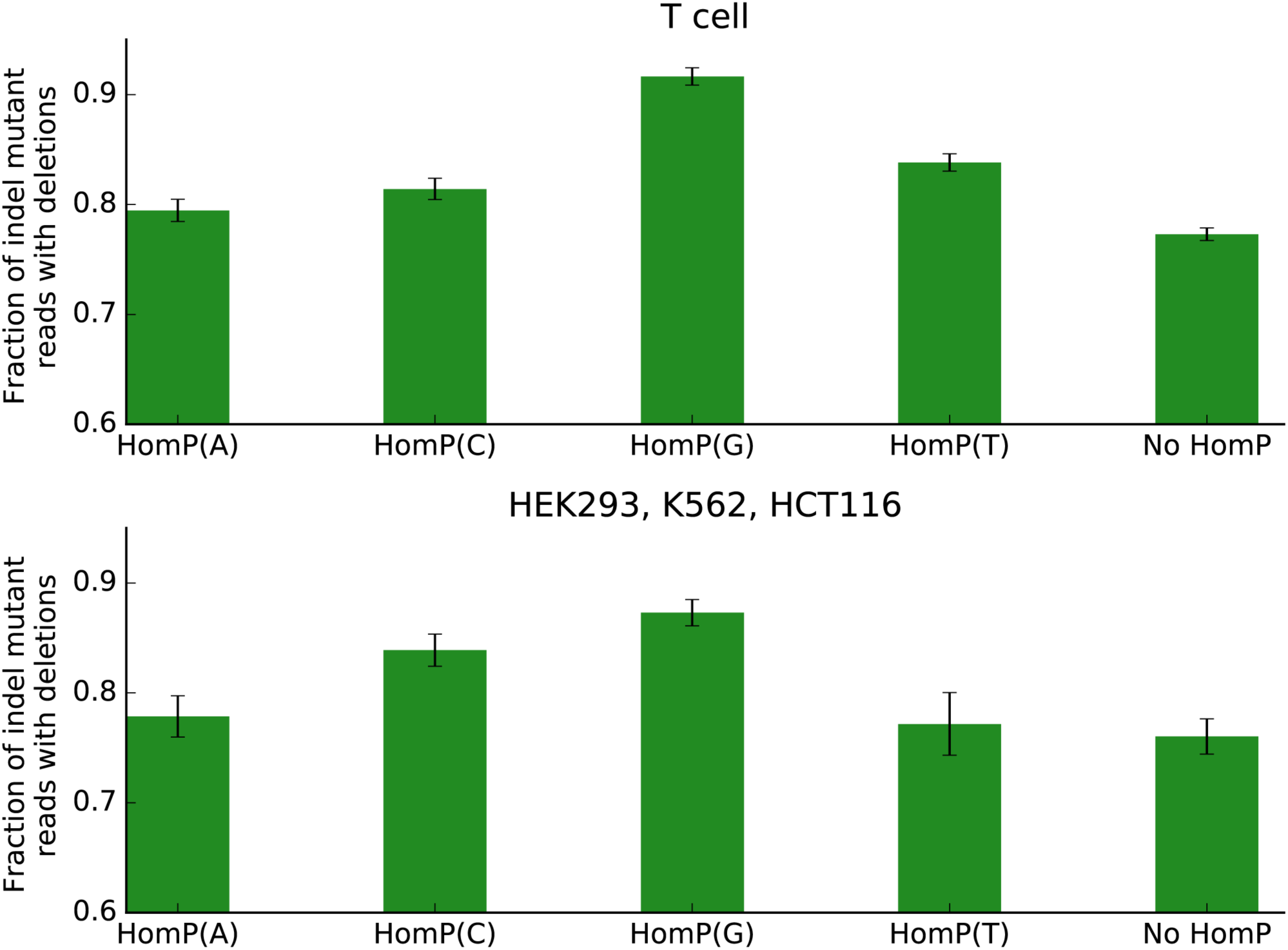
Average fraction of indel mutant reads with deletion in target sites grouped by their homopolymer types. HomP(A) corresponds to target sites that have at least two consecutive A nucleotides adjacent to the cut site, and similarly for HomP(C), HomP(G), and HomP(T). No HomP indicates the rest of the target sites without homopolymers. We show the results for T cells (top) and the aggregate results for HEK293, K562 and HCT116 (bottom). Error bars represent the standard error of the mean (SEM).

**Supplementary Figure 10.**
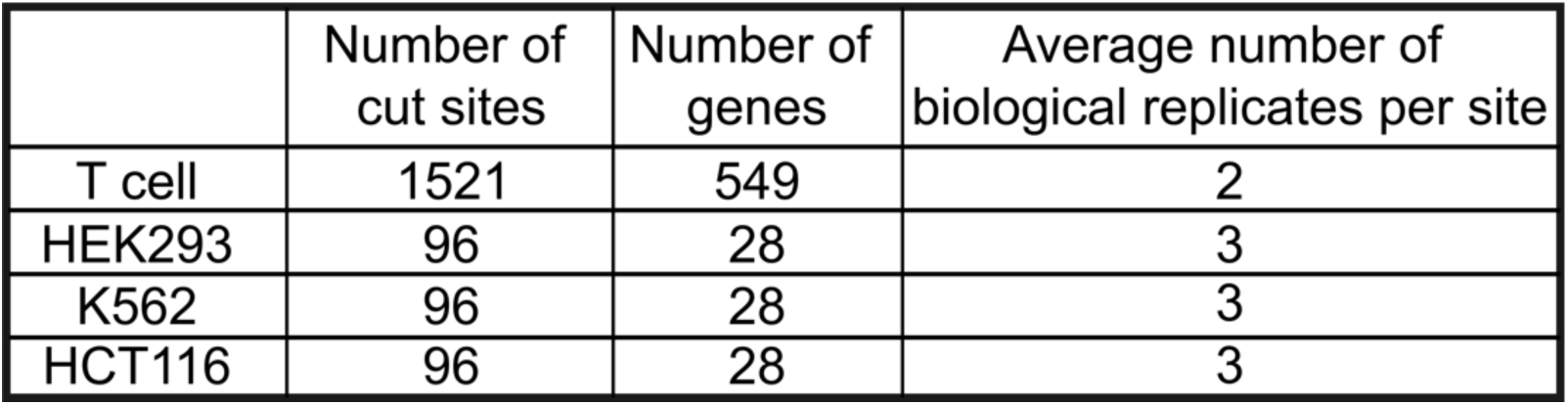
Data summary for T cells and three other cell types^5^.

**Supplementary Figure 11.**
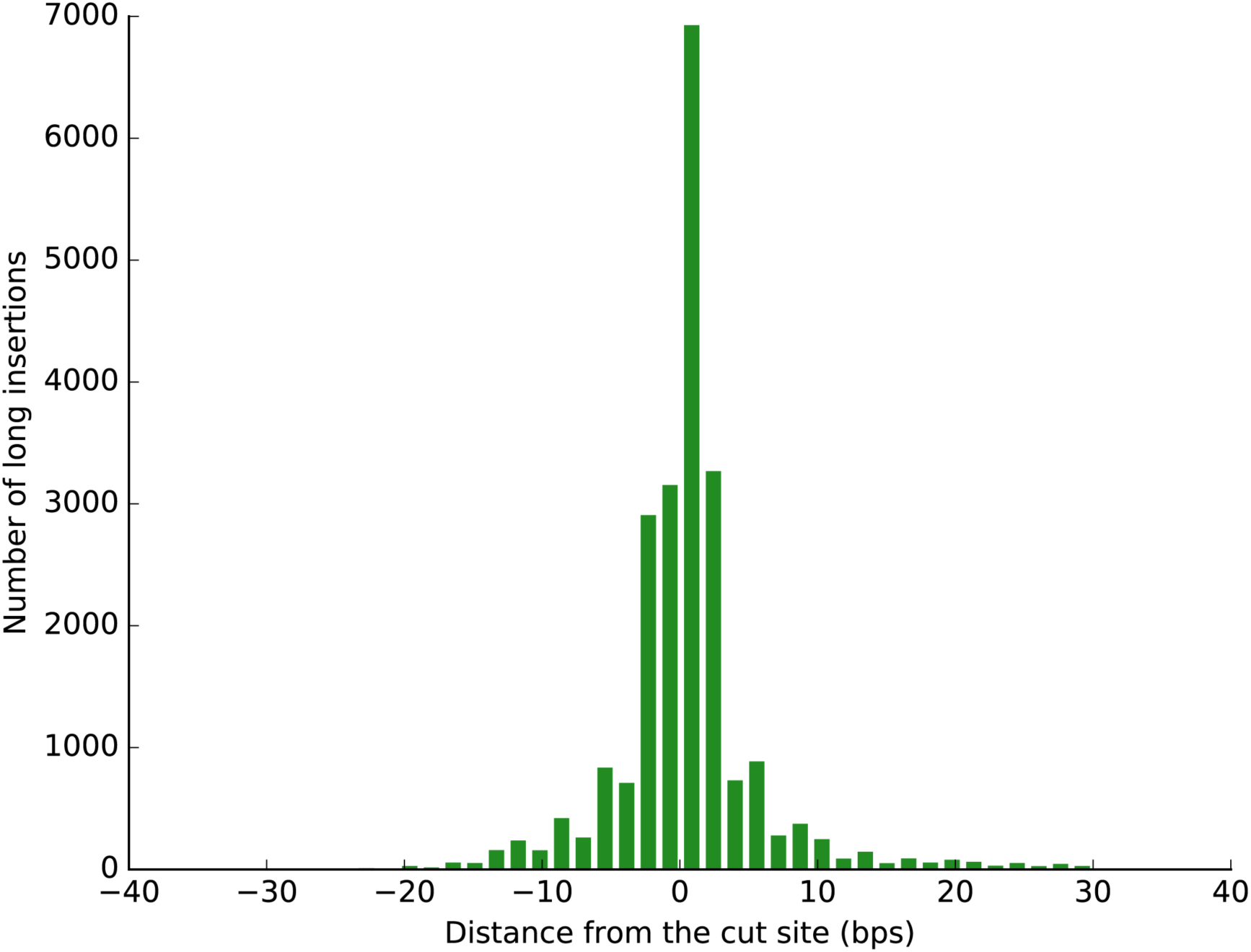
Histogram of the distances of long insertions from the target cut sites.

**Supplementary Figure 12.**
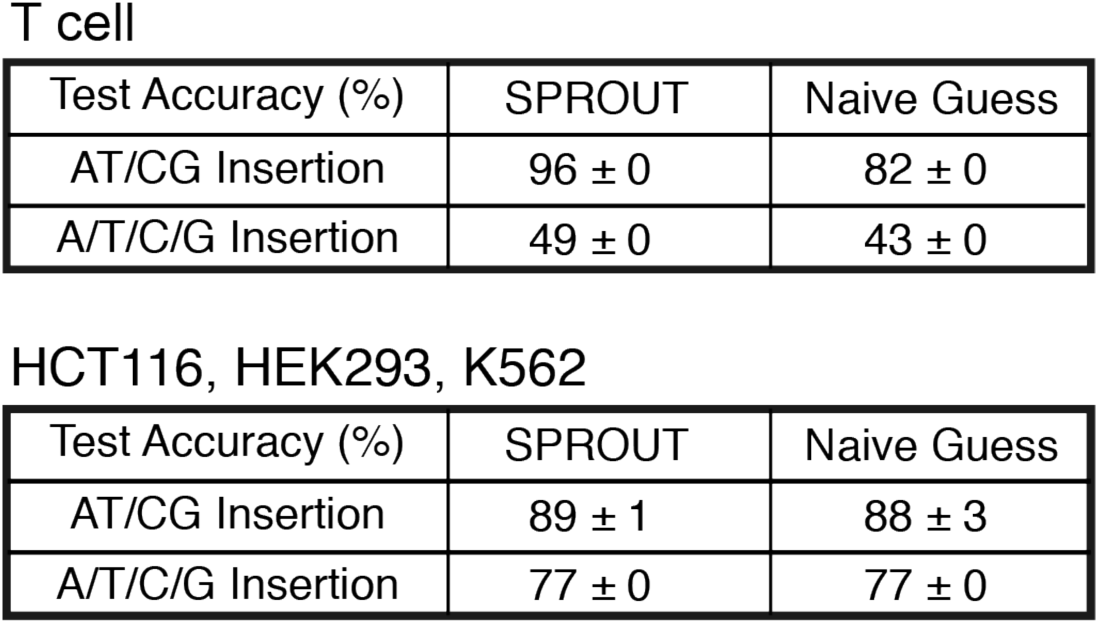
The performance of SPROUT in predicting the insertion type given a single-nucleotide insertion. The first row of each table gives the accuracy for predicting if the inserted nucleotide is A/T or C/G (binary classification). The second row of each table gives the accuracy of predicting if the inserted nucleotide is A, T, C, or G (4-class classification). SPROUT’s performance is compared to a naive guessing strategy which selects the most frequently occurring nucleotide(s) in the dataset.

**Supplementary Figure 13.**
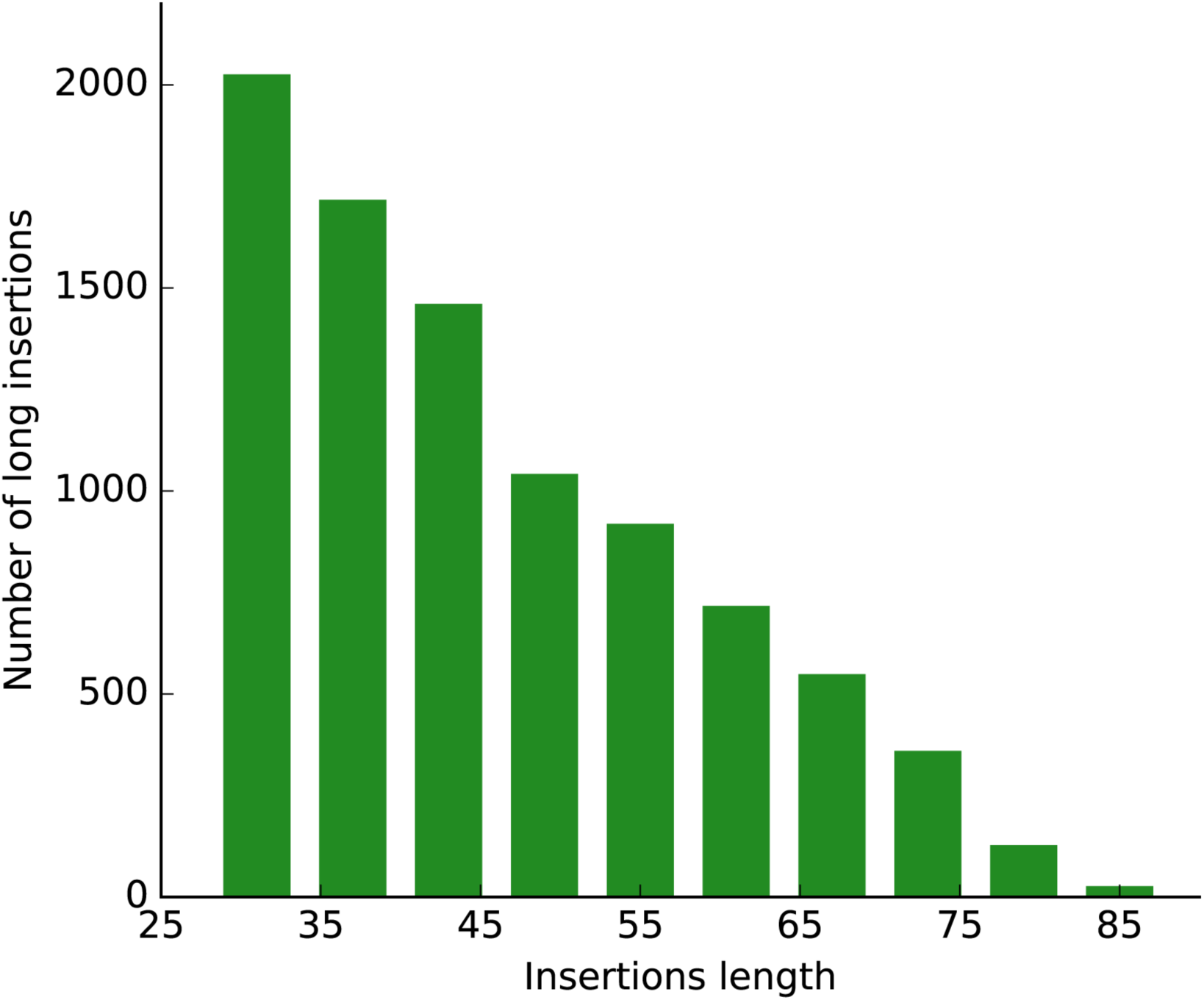
Distribution of the length of the aligned insertions in T cells.

**Supplementary Figure 14.**
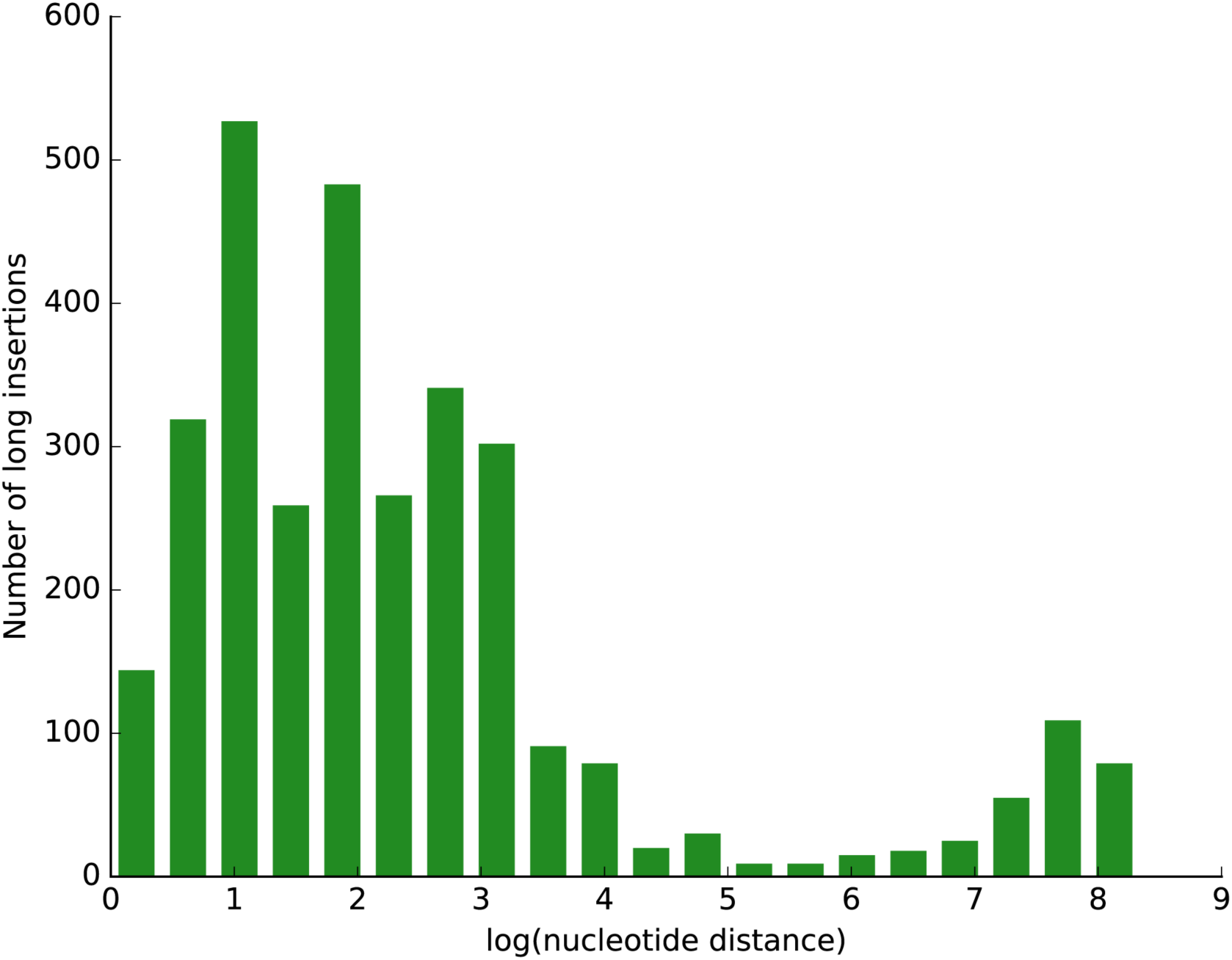
Histogram of the distance of the insertion donor sites to the cut sites in intra-chromosomal long insertions. The x-axis indicates distances in log 10 bases.

**Supplementary Figure 15.**
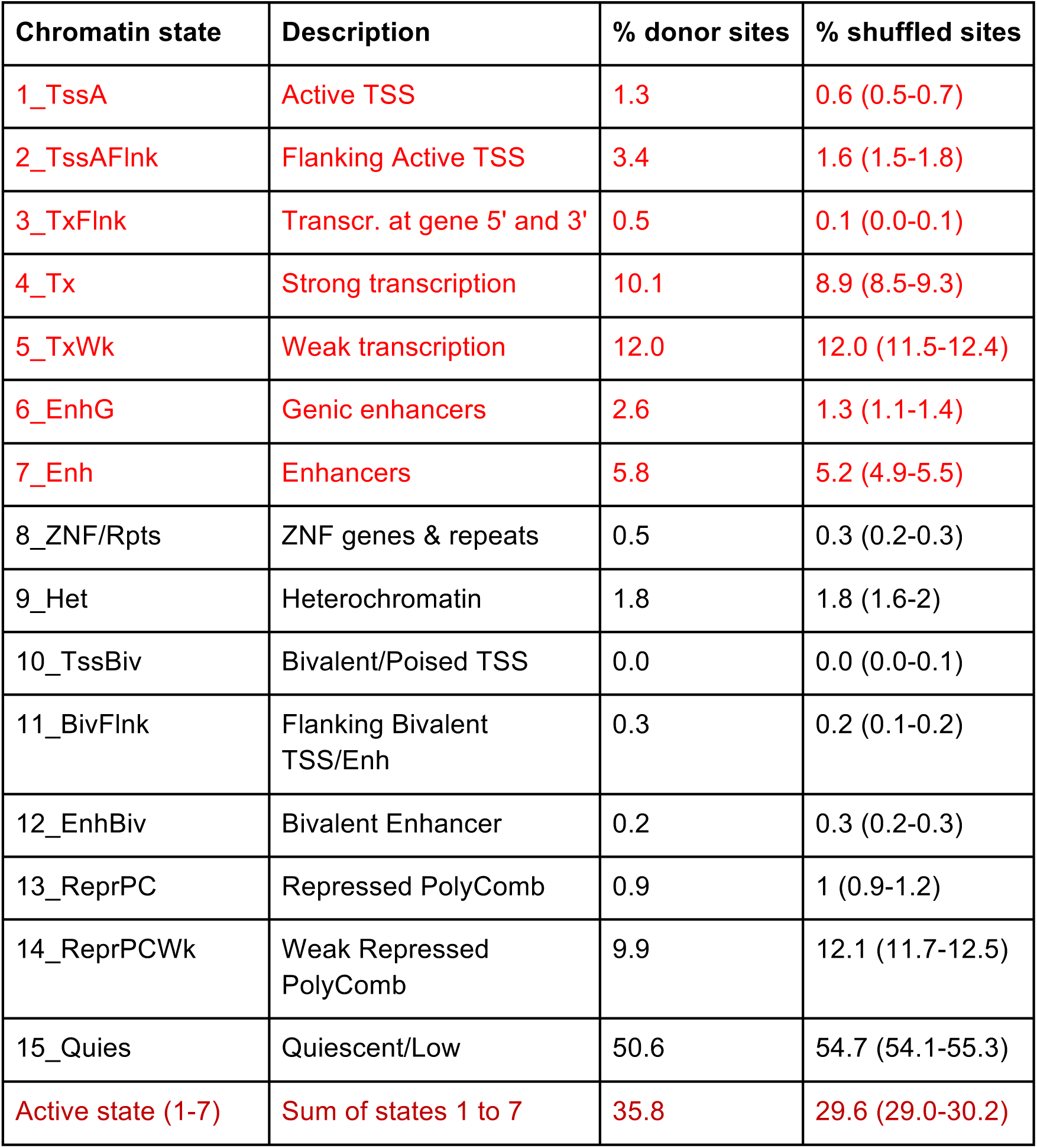
Overlap between the insertion donor sites and 15 core chromatin states. We measured the percentage of insertion donor sites that fall within each of the 15 chromatin states (“% donor sites”). The chromatin states were obtained for primary CD4+ T cells (E043) from the Human Epigenome Roadmap. Here we considered only the long insertions that are aligned to a *different* chromosome from the SpCas9 target site, to avoid potential confounding due the target sites being in exons. For background control, we randomly shuffled each aligned insertion within a +-500kb window centered at its original location (i.e. donor site), and report the percentage of the shuffled sites that overlap each chromatin state (“% shuffled sites”). Insertion donors are significantly enriched for chromatin states associated with enhancers and transcription (states 1-7, colored red) compared to control (P < 10^−5^). Altogether 35.8% of inter-chromosome donor sites come from one of states 1 to 7 compared to 29.6% of the control sites.

**Supplementary Figure 16.**
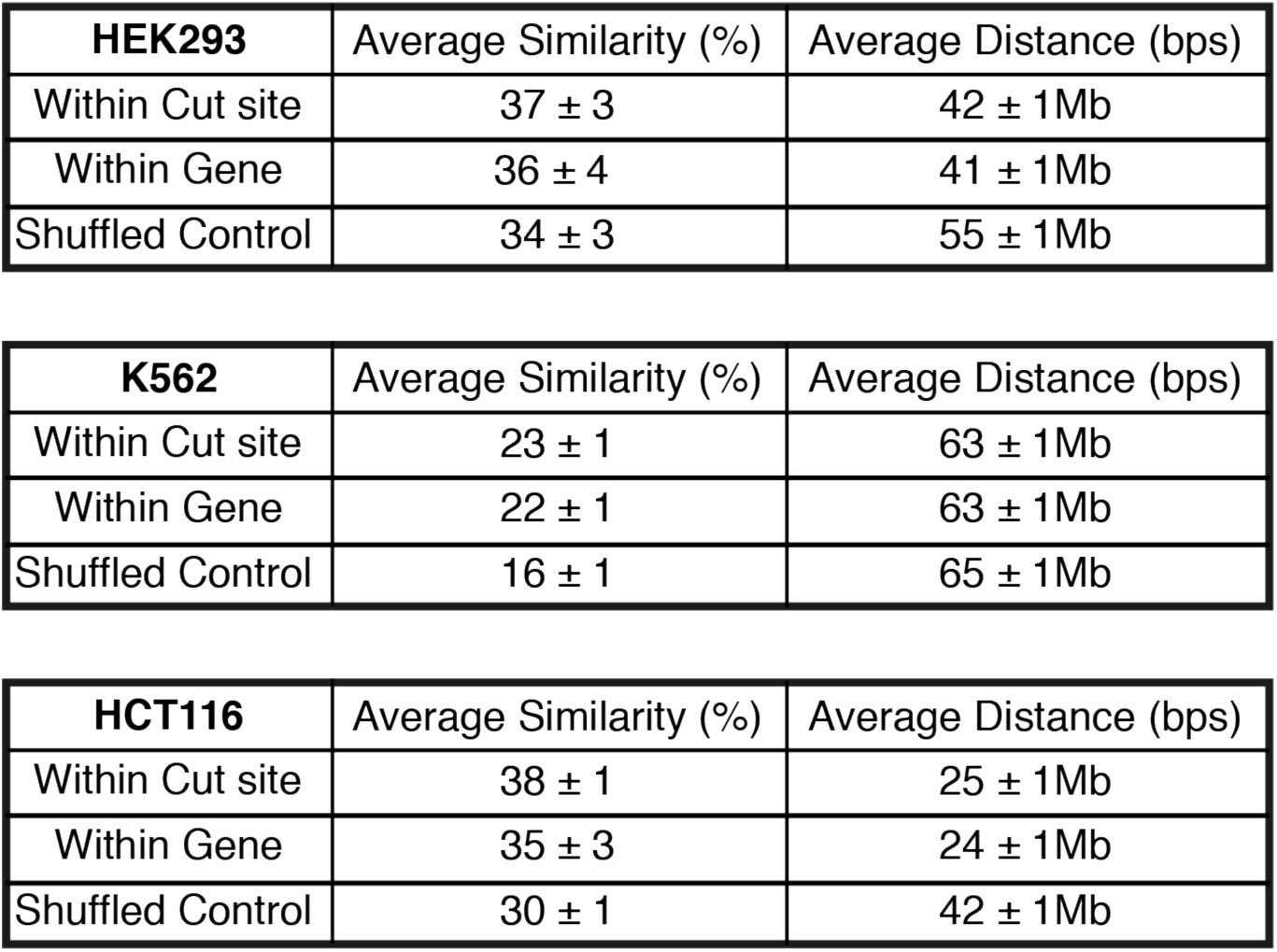
The average similarity and distance of the chromosomal positions between long insertions. We measure the similarity and distance for long insertions across biological samples at the same cut site (“Within cut site”), across different cut sites within the same gene (“Within gene”), and across random pairs of cut sites (“Shuffled control”). We report the results for each of three previously published data^5^.

**Supplementary Figure 17.**
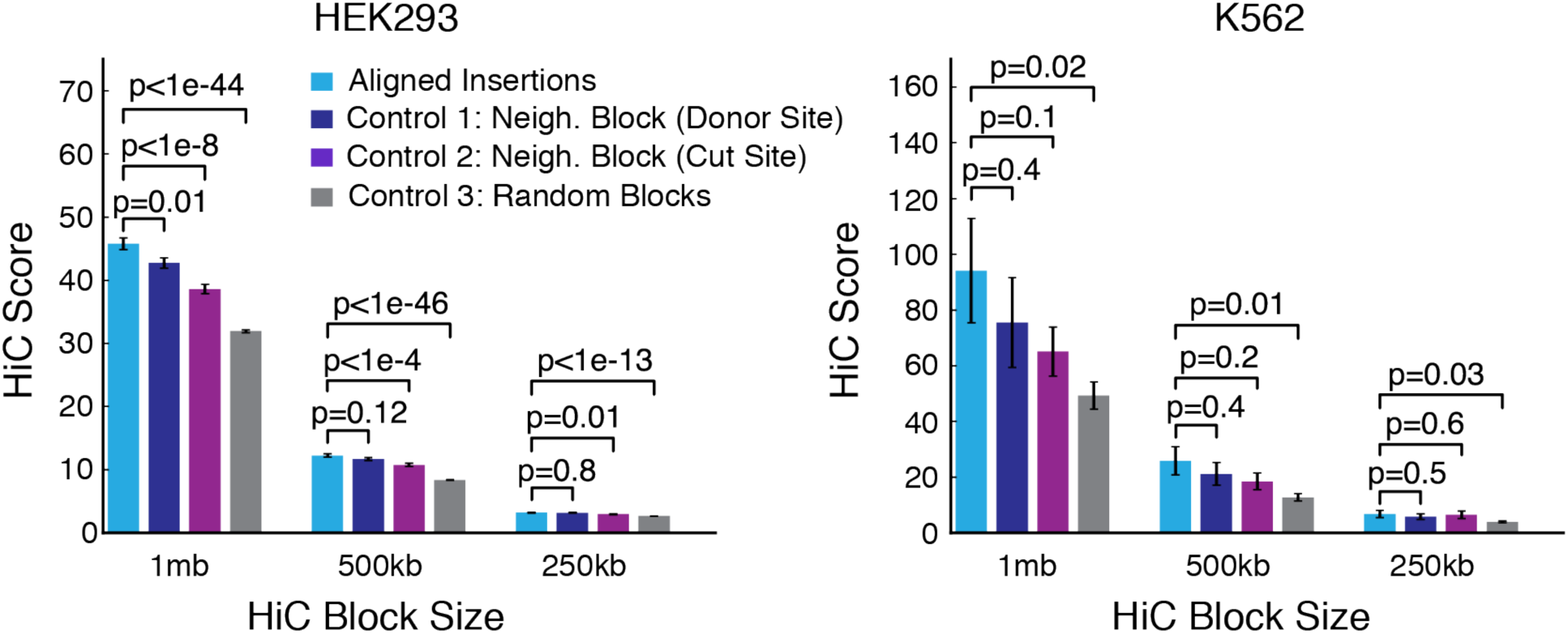
The HiC contact map at the insertion site locations compared to three control cases in two other cell types (HEK293 and K562) across different HiC block sizes for insertion larger than 25 nucleotides^5^. The first control averages the HiC contact map in the neighboring blocks of the insertion donor and cut site. The third control averages the HiC score among random blocks in the same cut site-donor site chromosome pairs. Error bars represent the standard error of the mean (SEM).

